# Structurally Distributed Surface Sites Tune Allosteric Regulation

**DOI:** 10.1101/2021.03.11.435042

**Authors:** James W. McCormick, Marielle A.X. Russo, Samuel Thompson, Aubrie Blevins, Kimberly A. Reynolds

## Abstract

Our ability to rationally optimize allosteric regulation is limited by incomplete knowledge of the mutations that tune allostery. Are these mutations few or abundant, structurally localized or distributed? To examine this, we conducted saturation mutagenesis of a synthetic allosteric switch in which Dihydrofolate reductase (DHFR) is regulated by a blue-light sensitive LOV2 domain. Using a high-throughput assay wherein DHFR catalytic activity is coupled to *E. coli* growth, we assessed the impact of 1548 viable DHFR single mutations on allostery. Despite most mutations being deleterious to activity, fewer than 5% of mutations had a statistically significant influence on allostery. Most allostery disrupting mutations were proximal to the LOV2 insertion site. In contrast, allostery enhancing mutations were structurally distributed and enriched on the protein surface. Combining several allostery enhancing mutations yielded near-additive improvements to dynamic range. Our results indicate a path towards optimizing allosteric function through variation at surface sites.

## INTRODUCTION

In allosteric regulation, protein activity is modulated by an input effector signal spatially removed from the active site. Allostery is a desirable engineering target because it can yield sensitive, reversible, and rapid control of protein activity in response to diverse inputs [1–3]. One common approach for achieving allosteric regulation in both engineered and evolved systems is through domain insertion: the transposition, recombination, or otherwise fusion of an “input” domain into an “output” domain of interest [4–7]. In natural proteins, domain insertions and rearrangements play a key role in generating regulatory diversity, with kinases serving as a prototypical example [8–11]. In engineered proteins, domain insertions have been used to generate fluorescent metabolite biosensors [7], sugar-regulated TEM-1 β-lactamase variants [12], and a myriad of light controlled proteins including kinases, ion channels, guanosine triphosphatases, guanine exchange factors, and Cas9 variants [5, 13–18]. In all cases, domain insertion provides a powerful means to confer new regulation in a modular fashion.

However, naïvely created domain insertion chimeras sometimes exhibit relatively modest allosteric dynamic range, with small observed differences in activity between the constitutive and activated states [19]. These fusions then require further optimization by either evolution or empirical mutagenesis, but general principles to guide this process are largely absent. Which mutations tune or improve an allosteric system? Because we lack comprehensive studies of allosteric mutational effects in either engineered or natural systems, it remains unclear whether such mutations are common or rare, and what magnitude of allosteric effect we might typically expect for single mutations. Additionally, it is not obvious if such mutations are structurally distributed or localized (for example, to the insertion site). Answers to these questions would inform practical strategies for optimizing engineered systems and provide insight into the evolution of natural multi-domain regulation in proteins.

To address these questions, we performed a deep mutational scan of a synthetic allosteric switch: a fusion between the *E. coli* metabolic enzyme Dihydrofolate Reductase (DHFR) and the blue-light sensing LOV2 domain from *A. sativa* [20, 21]. This modestly allosteric chimera shows a 28% increase in DHFR catalytic turnover (*k_cat_*) in response to light. Focusing on mutations to the DHFR residues, we found that only a small fraction (4.4%) of the mutations that retained DHFR activity had a statistically significant impact on allostery. Individual mutations exhibited generally modest effect sizes; the most allosteric single mutant characterized (H124Q) yielded a 3-fold increase in *k_cat_* in response to light relative to the starting construct. Structurally, allostery disrupting mutations tended to cluster near the LOV2 insertion site and were modestly enriched at both conserved and co-evolving amino acid positions. In contrast, allostery enhancing mutations were distributed across the protein, and strongly associated with the protein surface. We observed that combining a few of these mutations yielded near-additive enhancements to allosteric dynamic range. Collectively, our data elucidates practical strategies for optimizing engineered systems, and shows that weakly conserved, structurally distributed surface sites can contribute to allosteric tuning.

## RESULTS

### Characterization of an unoptimized allosteric fusion of DHFR-LOV2

To begin our study of allostery tuning mutations, we selected a previously characterized synthetic allosteric fusion between DHFR and LOV2 generated in earlier work [20, 21]. In this fusion, the LOV2 domain of *A. sativa* is inserted between residues 120 and 121 of the *E. coli* DHFR βF-βG loop; we refer to this construct as DL121 (**Figure 1A, B**). The choice of LOV2 insertion site was guided by Statistical Coupling Analysis (SCA), an approach for analyzing coevolution between pairs of amino acids across a homologous protein family [22–24]. A central finding of SCA is that co-evolving groups of amino acids, termed *sectors*, often form physically contiguous networks in the tertiary structure that link allosteric sites to active sites [24–26]. To create the DL121 fusion, Lee et al. followed the guiding principle that sector connected surface sites in DHFR might serve as preferred sites (or “hot spots”) for the introduction of allosteric regulation [20]. The resulting DL121 fusion covalently attaches the N- and C-termini of LOV2 into a sector connected surface on DHFR, and displays a two-fold increase in DHFR hydride transfer rate (*k_hyd_*) in response to blue light [20]. Under steady state conditions, we measured a 28% increase in the turnover number (*k_cat_*) in response to light and a statistically insignificant change in the Michaelis constant (K_m_) (**Figure 1C**). Thus, the DL121 fusion is modestly allosteric *in vitro*. As DHFR has no known natural allosteric regulation, the LOV2 insertion confers a new, evolutionarily unoptimized regulatory input.

**Figure 1.**
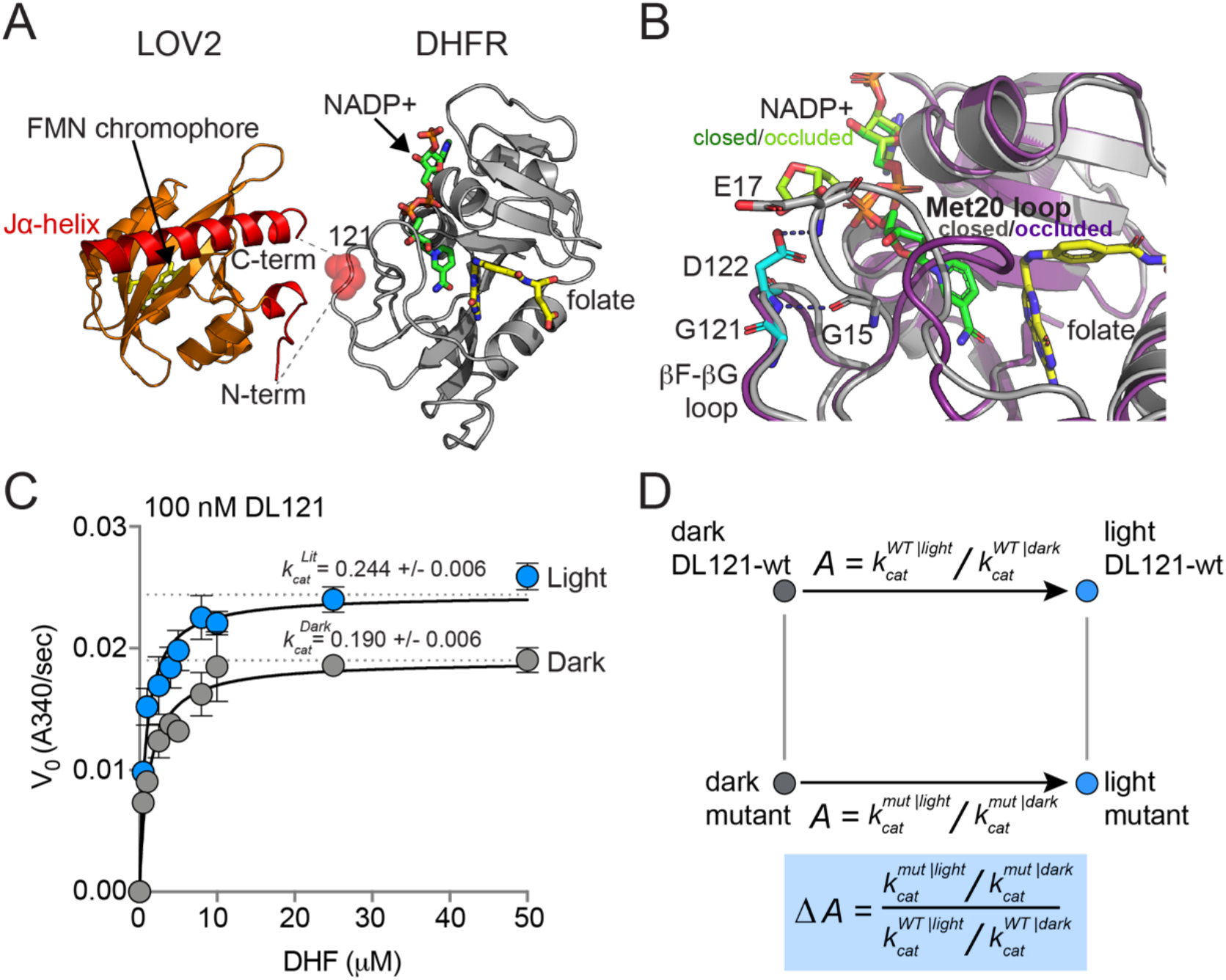
The DL121 DHFR/LOV2 fusion. **A**) Composite structures of the individual DHFR and LOV2 domains (PDB ID: 1RX2 and 2V0U), indicating the LOV2 insertion site between positions 120 and 121 of DHFR [28, 29]. DHFR is in grey cartoon, NADP co-factor in green sticks, and folate substrate in yellow sticks. In LOV2 signaling, blue light triggers the formation of a covalent adduct between a cysteine residue (C450) and a flavin mononucleotide (FMN, yellow sticks) [30–32] and associated unfolding of the C-terminal Jα-helix (red cartoon); this order-to-disorder transition is used for regulation in several synthetic and natural systems [33, 34]. **B**) DHFR loop conformational changes near the LOV2 insertion site. While the mechanism of DHFR regulation by LOV2 is currently unknown, inspecting the native DHFR structure provides some insight. The substrate-bound Michaelis complex of native DHFR is in the “closed” conformation (grey cartoon), while the product ternary complex is in the “occluded” state (purple cartoon). The βF-βG loop, where LOV2 is inserted, is highlighted in cyan. In native DHFR, hydrogen bonds between this loop (Asp122) and the Met20 loop (Gly15, Glu17) are thought to stabilize the closed conformation [28, 35]. Mutations to positions 121 and 122 reduce activity, and cause the enzyme to prefer the occluded conformation [36–38]. **C**) Steady state Michaelis Menten kinetics for the DL121 fusion under lit (blue) and dark (grey) conditions. The *k*_cat_ of DHFR increases 28% in response to light; the difference in K_m_ is statistically insignificant (Table S1). Error bars represent standard deviation for three replicates. **D**) Quantifying the allosteric effect of mutation. Allostery for the DL121 fusion is reported as the ratio between lit and dark catalytic activity. The effect of a mutation on allostery is then computed as the ratio of mutant allostery to wt-DL121 allostery (bottom blue box).

But can this relatively small allosteric effect generate measurable physiological differences that could provide the basis for evolutionary selection? DHFR catalyzes the reduction of 7,8-dihydrofolate (DHF) to 5,6,7,8-tetrahydrofolate (THF) using NADPH as a co-factor. THF then serves as a one-carbon donor and acceptor in the synthesis of thymidine, purine nucleotides, serine, glycine, and methionine. Because of these critical metabolic functions, DHFR activity is strongly linked to growth rate, and under appropriate conditions, *E. coli* growth rate can be used as a proxy for DHFR activity [21, 27]. Prior work found that the modest *in vitro* allosteric effect of DL121 conferred a selectable growth rate advantage *in vivo*: when an *E. coli* DHFR deletion strain (ER2566 *ΔfolAΔthyA*) was complemented with DL121, the resulting strain grew 17% faster in the light than in the dark [21]. Thus, DL121 is a system where: 1) allosteric control is rapidly and reversibly applied, 2) the allosteric effects on activity can be readily quantified both *in vitro* and *in vivo*, and 3) there remains potential for large improvements in regulatory dynamic range through mutation.

### A High Throughput Assay to Resolve Small Changes in DHFR Catalytic Efficiency

Our goal was to measure the effect of every single amino acid mutation in DHFR on the allosteric regulation of DL121. To do this, we aimed to follow a strategy loosely akin to a double mutant cycle (**Figure 1D**). The starting DL121 construct shows so-called V-type allostery, in which the effector (light) regulates the catalytic turnover number (*k_cat_*) [39]. Thus, allostery can be quantified as the ratio of catalytic turnover number between lit and dark states. More generally, allostery might be considered as a ratio of catalytic efficiencies (*k_cat_* /K_m_) between the lit and dark states, as the allosteric effector could regulate turnover, substrate affinity, or both. In either case, we defined the allosteric effect of mutation as the fold change in allosteric regulation upon mutation (**Figure 1D**, blue box). We sought to infer this quantity for every mutation in a saturation mutagenesis library of DHFR by using growth rate as a proxy for catalytic activity.

As in prior work, we measured the growth rate of many *E. coli* strains in parallel by using next generation sequencing (NGS) to monitor the frequency of individual DHFR mutants over time in a mixed culture (**Figure 2**) [21, 27]. Allele frequencies (*f_a_*) at each time point (t) were normalized as follows: 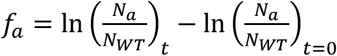 where *N_a_* and *N_WT_* are the number of mutant and wildtype (WT) counts at a given time point. By performing a linear fit of the log normalized allele frequencies vs. time, we calculated a slope corresponding to relative growth rate: this value is the difference in growth rate for the mutant relative to a reference (“WT”) construct.

**Figure 2.**
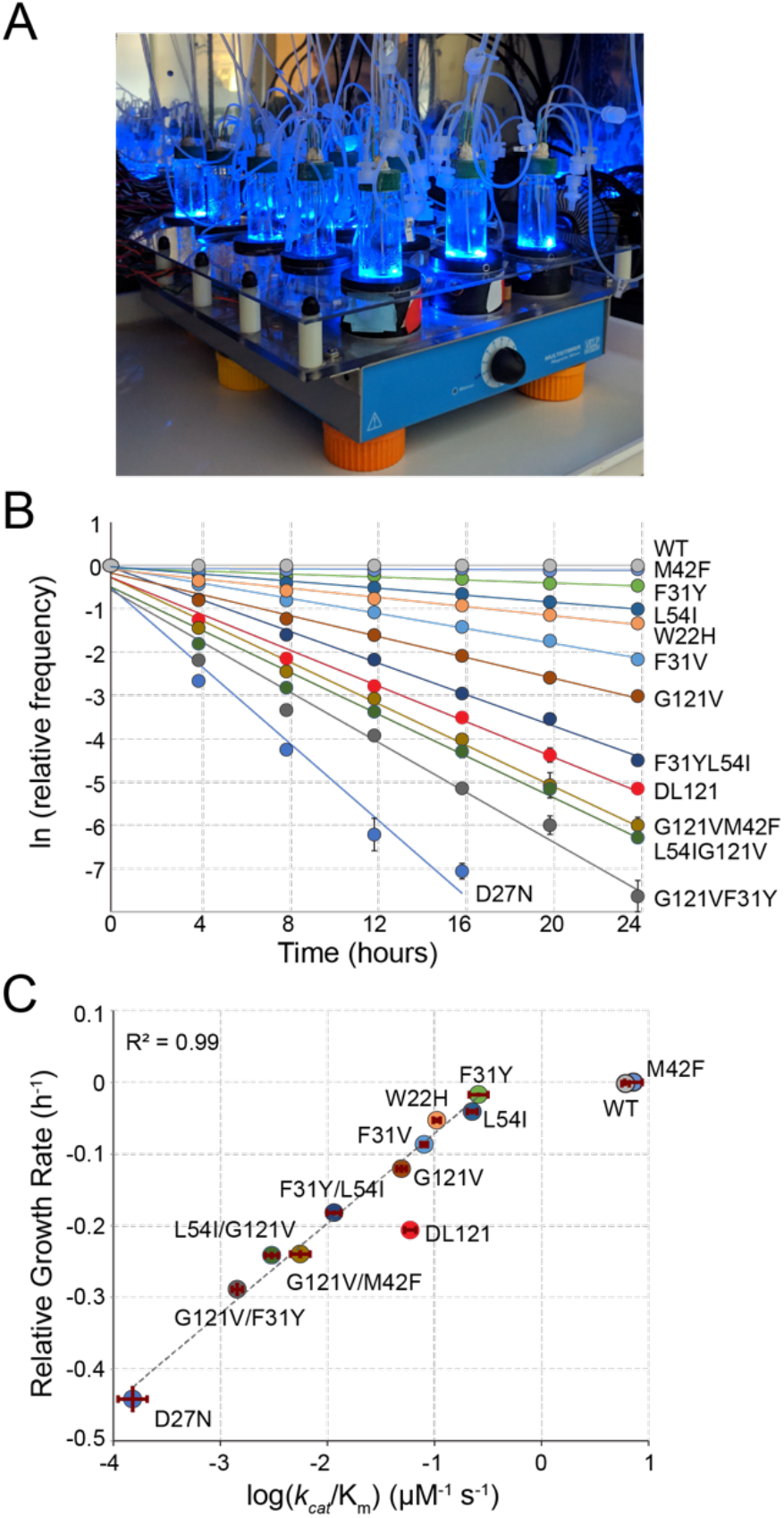
A high-throughput, high-resolution assay for DHFR activity. **A**) The turbidostat. The instrument has 15 individual growth chambers (vials), positioned on a stir plate inside an incubator. Illumination was provided by blue LEDs in each vial holder. **B**) Log-normalized relative allele frequency over time for 11 DHFR point mutations of known catalytic activity and the DL121 fusion. Allele frequency (colored circles) was determined by next generation sequencing of mixed-population culture samples at each time point. All frequencies were normalized to t = 0 and WT DHFR (no LOV2 insertion). Error bars reflect standard error across four measurements, they are sometimes obscured by the marker. The slope for each line of best fit provides the growth rate of each mutant allele relative to WT DHFR. **C**) Relative growth rate vs. log_10_(catalytic efficiency) for the 11 DHFR mutants and DL121 as characterized in panel B. Color coding of mutations is matched to panel B. Error bars reflect standard error of the mean over four replicates. The dashed line was fit by linear regression to all mutants in the linear regime (WT and M42F excluded).

As individual mutations tend to exhibit modest effects on allosteric regulation, we optimized the linear regime and resolution of the growth rate assay in two ways [21]. First, we grew the *E. coli* populations in a turbidostat outfitted with blue LEDs to activate LOV2 (**Figure 2A**). The turbidostat maintains each culture in exponential growth by dynamically sensing optical density and adjusting media dilution rate accordingly [40]; this ensures near-constant media conditions and eliminates the need for manual serial dilutions. Second, we selected media conditions – M9 minimal media with 0.4% glucose and 1 μg/ml thymidine supplementation – in which growth rate can resolve subtle differences in catalytic activity near the DL121 fusion. We evaluated the resolution of our assay using a “standard curve” of 11 point mutations of known catalytic activity in non-chimeric DHFR (**Figure 2B**). Under these conditions, we observed a log-linear relationship between relative growth rate and DHFR catalytic efficiency over nearly four orders of magnitude; this relationship saturates (plateaus) for the most active mutants (WT and M42F, **Figure 2C**). Importantly, the relative growth rate and catalytic activity of DL121 were near the center of the linear regime of our assay. We noticed that the DL121 chimera did not fall on the same “calibration line” as the point mutants of (non-chimeric) DHFR. Because variation in intracellular abundance also contributes to *in vivo* reaction velocity, one possibility is that DL121 was simply less abundant than the non-chimeric point mutants. In any case, because allostery concerns a relative change activity, light-independent differences in expression or abundance can be removed by appropriate normalization (as discussed further below).

As previously observed, the exponential divergence of mutants with different growth rates in a population makes it possible to detect even small biochemical effects [41]. More specifically, we can discriminate a change of +/- 0.02 μM^-1^ s^-^1 in catalytic power (*k_cat_*/K_m_) under these conditions. This level of precision is on par with – and in some cases better than – literature-reported errors for *in vitro* steady state kinetics measurements of DHFR [21, 42, 43]. Consequently, we can resolve small catalytic and allosteric effects of mutations on DL121 through this high-throughput growth-based assay.

### Deleterious Mutations are Enriched at Conserved, Coevolving Positions in DHFR

In order to map the coupling of individual DHFR positions to light, we constructed a deep mutational scanning library over all DHFR positions in the DL121 fusion (**Figure S1-2**). Then, we measured the growth rate effect of each mutation in triplicate under both lit and dark conditions using the above-described assay (**Figure 3A-C, Figure S3-4**). In this experiment, all growth rates were calculated relative to the unmutated DL121 fusion, which itself exhibits reduced activity (and growth rate) compared to WT DHFR. Mutations fell into four broad categories in terms of growth rate effects: neutral, uniformly deleterious (**Figure 3A**), uniformly beneficial (**Figure 3B**), or light dependent (and thus allosteric, **Figure 3C**). We were unable to measure growth rate for 891 of the 3021 possible missense mutations (19 substitutions over 159 positions): 226 (7.5%) were missing at the start of the experiment (t=0) for one or more replicates (referred to as “no data”), and an additional 665 (22%) were depleted from the library before reaching the minimum of three time points required for growth rate estimation (we refer to these as null mutants, see also methods, **Figure S4**). We interpreted these 665 rapidly depleting null mutants as highly deleterious to growth rate and thus DHFR activity. The relative growth rates for the remaining 2130 mutations (70.5%) were highly reproducible, with a correlation coefficient between replicate pairs above 0.9 (**Figure S3**).

**Figure 3.**
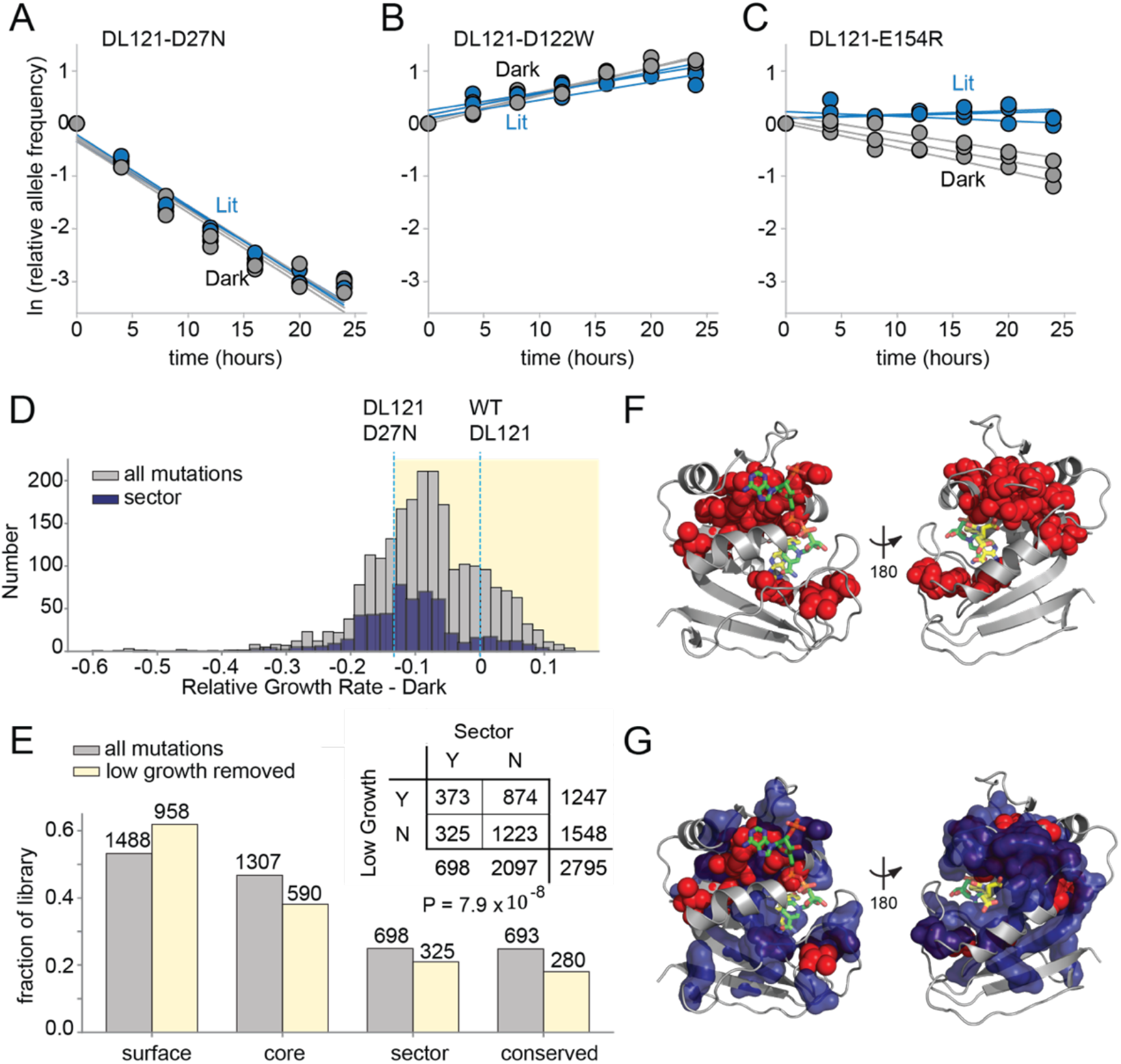
The effect of DL121 DHFR mutations on growth rate. **A-C**) Representative relative growth rate trajectories for three mutations. **A**) DL121 D27N was deleterious in both lit and dark conditions. **B**) DL121 D122W was advantageous under both lit and dark conditions. **C**) DL121 E154R was deleterious in the dark, and near neutral in the light. Solid lines were obtained by linear regression; the slope of these provides the difference in growth rate relative to the unmutated DL121 construct. Relative growth rates were measured in triplicate for each mutant under lit (blue) and dark (grey) conditions. **D**) Distribution of relative growth rates under dark conditions. The distribution for all mutations with measurable growth rate effects is in grey (‘null data’ and ‘no data’ excluded); the distribution for sector mutations is in navy. The relative growth rate of DL121 D27N, a mutation that severely disrupts catalytic activity, is indicated with a cyan dashed line. **E**) The fraction of DL121 mutations with measurable growth rates that can be categorized as: DHFR surface, core, sector, and evolutionarily conserved (see methods for definitions). The fraction is shown for both the complete library (grey bars), and the library after removing mutations with low growth (growth rate <= DL121 D27N). The absolute number of mutations is shown above each bar. A contingency table summarizes the overlap between mutations in the sector (at a p-value cutoff of 0.010), and the mutations that yield low growth (growth rate <= DL121 D27N). **F**) Structural distribution of positions enriched for mutations with growth rates as low as or lower than DL121 D27N (red spheres). The DHFR backbone is in grey cartoon, the folate substrate in yellow sticks, and the NADP co-factor in green sticks. **G**) Relationship of the sector (navy blue surface) to positions enriched for growth-rate disrupting mutations (red spheres, same as in C).

Before examining the allosteric effects of mutations, we first considered the effects of mutations on growth rate (and thus DHFR activity) in a single state. Prior work has found that deleterious mutations are enriched at evolutionarily conserved positions and within the protein sector [44]. The DHFR sector was defined by analyzing coevolution in a multiple sequence alignment of native DHFR domains, so we wished to examine if sector positions were indeed critical to function in the chimeric DL121 fusion. Good correspondence between the DHFR sector, evolutionary conservation, and deleterious mutations in DL121 would provide confidence that the core functional elements of native DHFR remain intact in the chimera. The vast majority of mutations were at least modestly deleterious to growth, with a median relative growth rate of – 0.084 in the dark and −0.083 in the light (**Figure 3D**). A cluster of beneficial mutations was observed just before the LOV2 insertion site at position 121 in both conditions, suggesting some potential to compensate for the inserted LOV2 (**Figure S4**). The overall distribution of fitness effects shows some differences relative to prior DMS studies of natural proteins including native *E. coli* DHFR [27, 45]. First, the distribution of fitness effects for mutations in natural proteins is often centered around neutral, implying a certain degree of mutational robustness [44, 46]. Secondly, DMS of native DHFR – under experimental conditions designed to resolve mutational effects near WT – revealed many beneficial (activating) mutations [27]. There are two explanations for the relative paucity of beneficial and neutral mutations in the present dataset. First, the DL121 fusion is comparably less robust because the unoptimized LOV2 insertion introduces an initial compromise to DHFR function. Secondly, the conditions of our assay (both expression and media) differ from prior work [27] and were selected to resolve mutational effects near DL121; consequently, mutations with native-like (or better) activity are in the saturating, nonlinear regime of our assay.

To identify the slowest growing – and presumably near, or entirely, inactivating – mutations, we applied an empirical growth rate cutoff of −0.13 to the lit and dark growth rates. This corresponds to the growth rate for DL121 D27N; D27N is an active site mutation that strongly reduces the activity of WT DHFR (**Figure 2B, C**). The DL121 D27N mutant grows very slowly in the conditions of our assay and is inviable in the absence of thymidine supplementation (**Figure S5**). We found that mutations with growth rates at or below this cutoff (including the null mutants) were significantly enriched in both the sector (P=7.9×10^-8^, **Figure 3E**) and at evolutionarily conserved positions (P=8.7×10^-20^, **Figure S6**). When mapped to the WT DHFR structure, positions enriched for deleterious mutations surround the active site and co-factor binding pocket (**Figure 3F**), structurally overlap with the sector (**Figure 3G**), and include a number of positions known to play a critical role in WT DHFR catalysis (e.g. W22, D27, M42, and L54) [47, 48]. These data are consistent with the view that sector positions continue to play a key role in conferring DHFR catalytic activity in the DL121 fusion.

Following the thinking that (near) inactive DHFR variants are both inherently non-allosteric and associated with the least reproducible growth rate measurements (**Figure S3**), we removed the set of 1247 slow-growing (growth rate < −0.13) and null mutations prior to the analysis of allostery. The retained 1548 mutations – representing 51% of the growth assay data – remain well-distributed between the DL121 surface, core, sector, and evolutionarily conserved positions (**Figure 3E**). These present a high-confidence and representative subset of the data for evaluating mutational effects on DL121 allosteric regulation.

### Allostery Tuning Mutations are Sparse

To compute the allosteric effect of mutation, we considered the triplicate measurements of lit and dark relative growth rate for each mutant (**Figure 3A-C**). Given the log-linear relationship between growth rate and catalytic efficiency, subtracting growth rates approximates log-ratios of catalytic efficiencies. Thus, we estimated the allosteric effect of mutation by taking the difference in the average relative growth rates between lit and dark conditions:

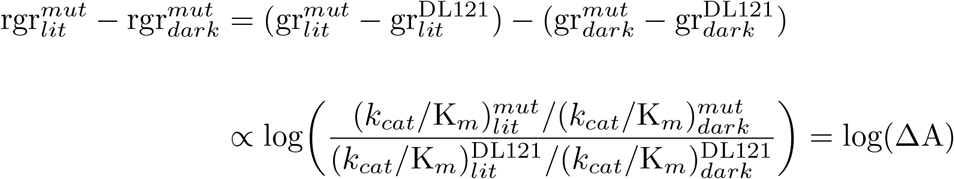

In the above equations, rgr is relative growth rate (which is directly measured in our sequencing-based assay) and gr refers to absolute growth rate. Accordingly, positive values indicate allostery enhancing mutations and negative values indicate allostery disrupting mutations (**Figure 1D, 4A**). Of the 1548 mutations evaluated, the allosteric effect is normally distributed with a mean near zero (μ = 0.0017, **Figure S7**). To assess the statistical significance of allosteric effects, we computed a p-value for each mutation by unequal variance t-test under the null hypothesis that the lit and dark replicate measurements have equal means. These p-values were compared to a multiple-hypothesis testing adjusted p-value of P=0.016 determined by Sequential Goodness of Fit (SGoF, **Figure 4A**) [49]. Under these criteria, only 69 mutations (4.5% of all viable mutants) significantly influenced allostery: 56 mutations enhanced allostery while 13 disrupted regulation.

**Figure 4.**
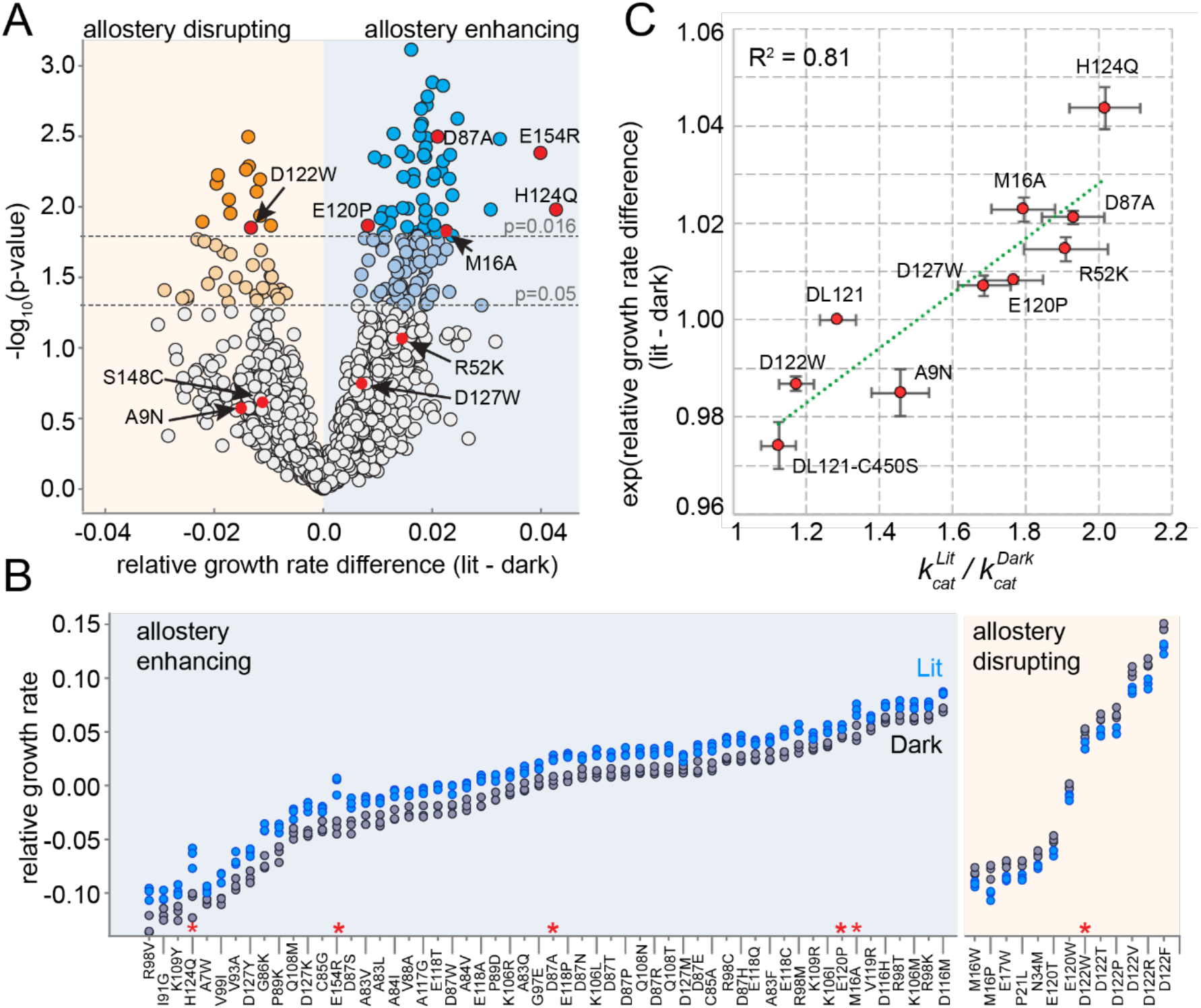
The effect of DL121 DHFR mutations on allostery. **A**) Volcano plot indicating the statistical significance of the light-dark growth rate difference (y-axis) as a function of relative growth rate difference (x-axis). P-values were computed using a t-test across triplicate light and dark measurements. Individual points correspond to mutations; mutations on the left (yellow) side of the graph are allostery disrupting, while mutations on the right (blue) are allostery enhancing. Two cutoffs for statistical significance are indicated with dashed grey lines – both a standard value of p = 0.05, and an adjusted p-value of 0.016, obtained by using Sequential Goodness of Fit (SGoF) to account for multiple hypothesis testing. Mutations selected for further *in vitro* experimental characterization are colored red and labeled. **B**) Triplicate relative growth rate measurements under lit (blue) and dark (grey) conditions for all mutations with statistically significant allostery at the adjusted p-value (p<=0.016). The mutations are sorted by dark growth rate; mutations selected for *in vitro* characterization are marked with red asterisks. **C**) Relationship between the allosteric effect as measured *in vivo* and *in vitro*. As we expect a log-linear relationship, we compare the ratio of catalytic constants (along x) to the exponent of the relative growth rate difference (along y). The relative growth rate difference under lit and dark conditions is the mean of triplicate measurements, error bars indicate SEM. All mutant effects on growth rate were measured in the same experiment (corresponding to a subset of the data in panel B) with the exception of DL121 C450S. The relative growth rate for this light-insensitive LOV2 mutant was measured in the ‘calibration curve’ experiment shown in figure 2 (see also methods). The ratio between *k*_cat_ in the light and *k*_cat_ in the dark reflects the mean of triplicate measurements; error bars indicate SEM. The green line was fit by linear regression.

We did not observe a strong association between the magnitude of growth rate effect and the allosteric effect size. Allostery-influencing mutations spanned a wide range of growth rates and exhibited comparatively modest effects on light regulation (**Figure 4B**).

To further examine the ability of the growth-based sequencing assay to quantitatively resolve mutation-associated changes in allosteric regulation, we selected 10 mutations spanning a range of allosteric and growth rate effects for *in vitro* characterization (**Figure 4A** red dots, **Figure S8-10**). As a control, we included the light insensitive variant DL121-C450S: the C450S mutation of LOV2 abrogates light-based signaling by blocking formation of a light induced covalent bond between position 450 and the FMN chromophore [50]. We expressed and purified the selected DL121 mutants to near homogeneity; S148C and E154R did not yield sufficient quantities of active protein for *in vitro* studies. For the remaining eight mutations we measured the *k_cat_* and K_m_ of DHFR under lit and dark conditions (**Figure S8**). To confirm function of the fused LOV2 domain, we also measured relaxation of the FMN chromophore following light stimulation and collected absorbance spectra before and after the application of light (**Figure S9, S10**). As expected, all of the characterized DL121 mutations (with the exception of DL121-C450S) retained LOV2 domains with light-responsive absorbance spectra and chromophore relaxation constants similar to the unmutated DL121 construct. Evaluating the light dependence of DHFR activity, the change in K_m_ value between lit and dark conditions was neither significant for any point mutation nor correlated to allosteric effect size (R^2^=0.003) (**Table S1**, **Figure S11A-C**). The K_m_ values for all characterized mutants (0.15-1.9 μM) were similar to that of unmutated DL121 (~1 μM). Given that the *in vivo* concentration of DHF in wildtype *E. coli* is well above the measured K_m_ (~25 μM [51]), we thought our assay might report more strongly on changes in *k_cat_*. Indeed, the values of *k_cat_* were more often significantly influenced by light (**Figure S11D**). The ratio of *k_cat_* in the light relative to the dark ranged from 1.12 (for the non-allosteric DL121-C450S construct) to 2.02 (for the most allosteric point mutation, H124Q) (**Figure 4B**). For reference, the starting DL121 construct has a lit:dark *k_cat_* ratio of 1.28. A comparison of the *in vitro* allosteric effect on *k_cat_*to the *in vivo* growth rate effect yields a near-linear relationship with a correlation coefficient of 0.81 (**Figure 4C, Figure S11D**). Taken together, these data show that our growth-based assay is quantitatively reporting on changes in allostery, and that the allosteric mutations identified here modulate DHFR activity through changes in catalytic turnover number.

### The Structural Pattern of Allostery Tuning Mutations

Next, we examined the distribution of allostery-tuning mutations on the WT DHFR tertiary structure. The 13 allostery disrupting mutations localized to six DHFR positions concentrated near the LOV2 insertion site (**Figure 5A**). More specifically, 90% of the allostery disrupting mutations occurred within 10Å of the DHFR 121 cα atom (**Figure 5B**). These mutations were modestly enriched in the protein sector (**Table S4**). Overall, the observed spatial distribution suggests these mutations may disrupt allostery by altering local structural contacts needed to ensure communication between DHFR and LOV2.

In contrast to this localized pattern, the 56 allostery enhancing mutations were observed at 25 positions distributed across the DHFR structure (**Figure 5C**) and enriched on the protein surface (**Figure 5D, Table S5**). These enhancing mutations were never found in the protein sector and were thus statistically significantly depleted from the protein sector (**Figure 5E, F**). This relationship – wherein allostery disrupting mutations were modestly enriched and allostery enhancing mutations were strongly depleted from the sector – also holds when defining the set of allosteric mutations at a relaxed cutoff of P=0.05 (**Table S4**). Given the prior finding that *sector connected* surface sites were hotspots for introducing allostery in DHFR [21], we also examined the association between allostery-influencing mutations and two other groups of DHFR positions: 1) surface sites that are either within or contacting the sector, and 2) surface sites that are only contacting the sector (but not within-sector). As for the analysis of sector positions only, we observed a statistically significant depletion of allostery enhancing mutations and enrichment of allostery disrupting mutations when considering the set of surface sites within or contacting the sector. This finding holds true over a range of significance thresholds for defining sector and allosteric mutations (**Table S6**). When considering the set of positions that contact (but are not within) the sector, we did not observe a statistically significant association at nearly all cutoffs (**Table S7**). Indeed, a number of allostery enhancing mutations do not contact the sector at all and occur in surface exposed loops (e.g. from residues 84-89, and from 116-119). So, counter to our expectations, the optimization of allostery did not occur at sector connected sites or even proximal to the LOV2 insertion site. Instead, structurally distributed and weakly conserved surface sites provided a basis for tuning and enhancing allosteric regulation regardless of sector connectivity.

**Figure 5.**
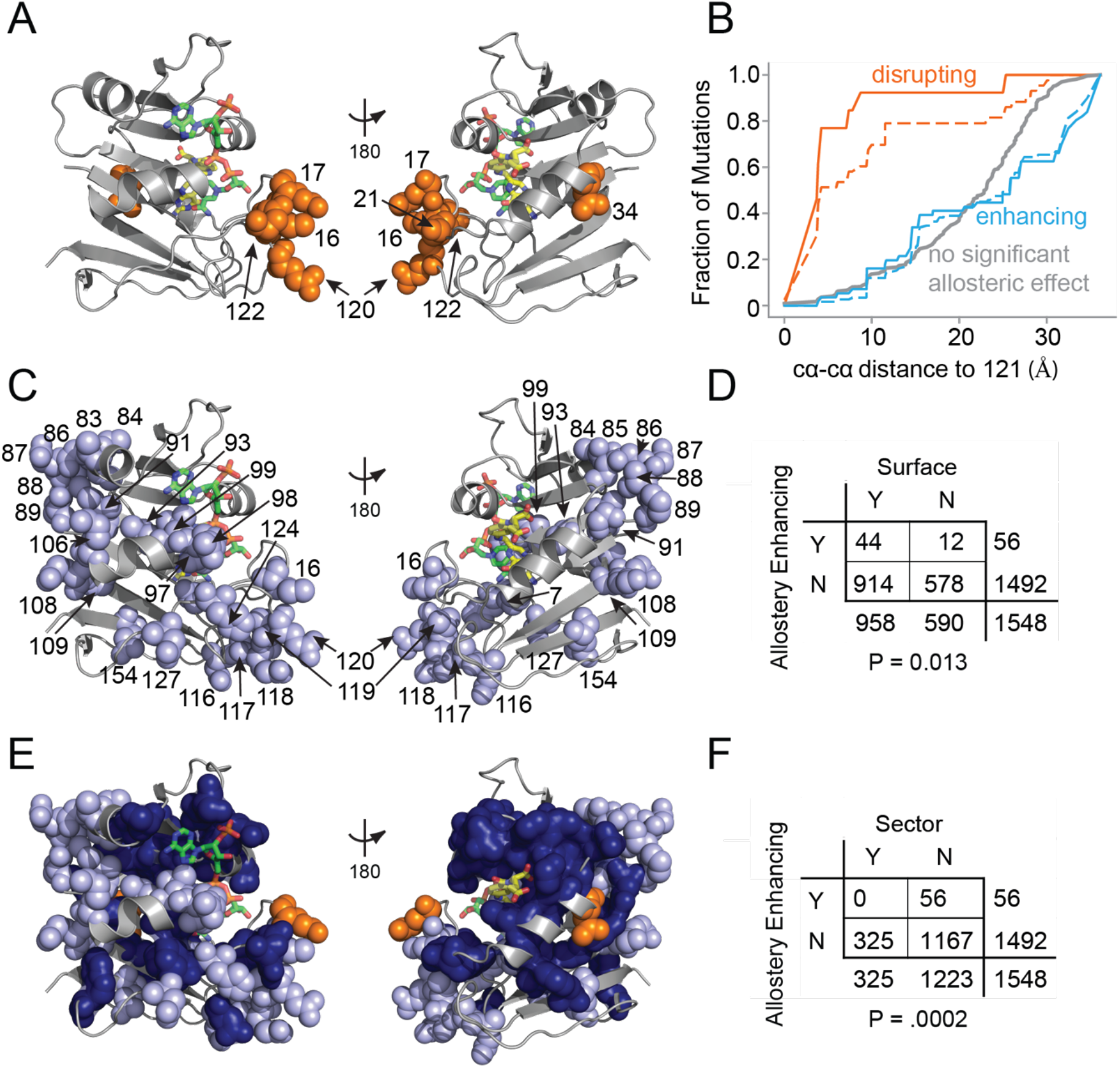
Structural distribution of allosteric mutations. **A**) Sites of allostery disrupting mutations (orange spheres). DHFR backbone is in grey cartoon, folate substrate in yellow sticks, and NADP co-factor in green sticks. **B**) Fraction of mutations that enhance (blue), disrupt (orange), or do not significantly influence allostery (grey) as a function of distance to the LOV2 insertion site at DHFR position 121. Solid and dashed lines indicate mutations at either the P=0.016 and P=0.05 significance cutoffs for allostery respectively. **C**) Sites of allostery enhancing mutations (light blue spheres). **D**) Contingency table summarizing the overlap between allostery enhancing mutations and mutations on the DHFR solvent accessible surface (considered as >25% relative solvent accessibility in the 1RX2 PDB). **E**) Sites of allostery enhancing (light blue spheres) and disrupting mutations (orange spheres) in the context of the sector (dark blue surface). **F**) Contingency table summarizing the relationship between allostery enhancing mutations and sector mutations (sector defined at a p-value cutoff of 0.010). No allostery enhancing mutations occur within the sector.

Taken together, our data show that many distributed surface sites can make modest contributions to allosteric regulation. Can these mutants be combined to further improve allosteric dynamic range? To test this, we created two mutant constructs by combining the most potent allostery enhancing mutations as characterized *in vitro*: the double mutant DL121-M16A,H124Q, and the triple mutant DL121-M16A,D87A,H124Q (**Figure 6A**). For both constructs we measured steady state catalytic parameters (**Table S1**) and verified LOV2 function through absorbance spectra and chromophore relaxation kinetics experiments (**Figure S12**). Interestingly, all three mutations exhibited near-log-additive improvements in allostery (**Figure 6B**). The DL121-M16A,H124Q fusion exhibits a 3.2 fold increase in *k_cat_* upon light activation while the triple mutant shows a 3.9 fold increase. For both mutant combinations, the improvement in allostery is realized by reducing the dark state (constitutive) activity (**Figure S12, Table S1**). The serial addition of allostery enhancing mutations reduced the overall catalytic activity of DHFR, suggesting that further improvement could be obtained by combining these mutations with a non-allosteric but activity-enhancing mutation. Overall, these data suggest that a naïve sector connected fusion can be gradually evolved towards increased allosteric dynamic range through the stepwise accumulation of single mutations at structurally distributed surface sites (**Figure 6C**).

**Figure 6.**
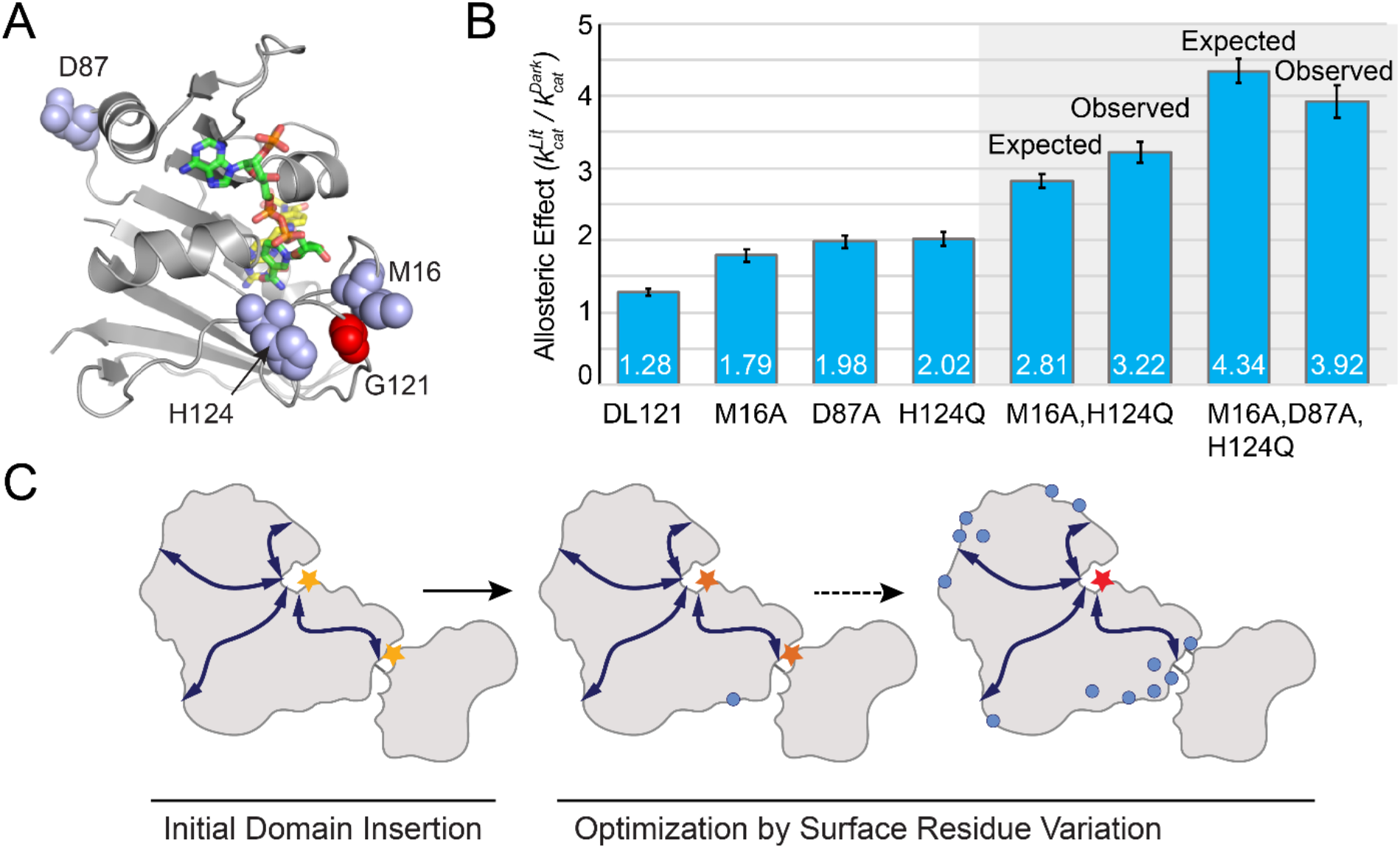
Combinatorial effect of allostery-enhancing mutations. **A**) Location of M16, D87 and H124 (blue spheres). The LOV2 insertion site, G121, is shown in red spheres. The DHFR backbone is in grey cartoon, the folate substrate in yellow sticks, and the NADP co-factor in green sticks. **B**) The *in vitro* allosteric effect of the single, double and triple mutants. Included are the log-additive expectations (Expected) for the double and triple mutants as well as the experimentally measured effects (Observed). The ratio between *k*_cat_ in the light and *k*_cat_ in the dark reflects the mean of triplicate measurements; error bars indicate SEM. **C**) Schematic whereby a novel domain insertion is iteratively optimized by surface residue variation.

## DISCUSSION

We used deep mutational scanning to study the frequency and structural pattern of allostery tuning mutations in a synthetic allosteric system, with the goal of understanding how regulation between domains can be optimized. Overall, allostery-influencing mutations were rare – just under 5% of viable mutations had statistically distinguishable effects on the lit and dark states of the DL121 fusion. We found that mutations at conserved and co-evolving (sector) positions were often deleterious to DHFR function and infrequently influenced allosteric regulation. In a few cases, sector mutations served to disrupt allostery; and nearly all allostery disrupting mutations were localized to the LOV2 insertion site on DHFR. Counter to our expectations, allostery enhancing mutations were distributed across the DHFR structure, depleted from the sector, and enriched on the protein surface. When considered individually, the allostery-enhancing mutations had modest effects (up to 2-fold) on regulation, but (at least in some cases) they can be combined to yield near-additive improvements in dynamic range. A triple mutant (DL121-M16A,D87A,H124Q) rationally designed using our point mutant data produces a 3.9 fold increase in *k_cat_* upon light stimulation, up from the 1.28 fold allosteric effect of our starting construct.

These results should be considered in the context of our experiment: the DL121 fusion begins with sharply reduced DHFR activity, and our experiment intentionally used relatively stringent DHFR selection conditions to better resolve small biochemical differences. Thus, it is unsurprising that a large fraction of DHFR mutations in our library were deleterious, with an appreciable fraction near-inactive. This result echoes prior studies showing that the fraction of deleterious mutations (and mutational robustness) is strongly modulated by a variety of factors, including purifying selection strength and expression level [46, 52, 53]. Given the finding that stabilizing mutations can often improve protein evolvability [53–55], it would be interesting to examine how the distribution of mutational effects on both DL121 function and allostery would change in the background of a stability (and/or activity) enhancing mutation to DL121. Nevertheless, our data serve to illuminate the pattern of mutational effects on a newly established (and unoptimized) domain fusion – the presumptive first step towards regulation in a number of both natural and synthetic systems.

Interestingly, we observe a seeming disparity between the sites where we were able to introduce new allosteric regulation by domain fusion (in our earlier work), and the sites where allosteric tuning takes place (in this work). Previously, Reynolds et al. found that sector connected surface sites served as hotspots for the introduction of new light-based regulation in DHFR [21]. Indeed, allosteric regulation was never obtained when the LOV2 domain was inserted at a non-sector connected site. In contrast, in this work we observed that allostery enhancing mutations were depleted both within the sector and at sector connected sites. For example, we observed a number of allostery enhancing mutations at positions 83-89 of the DHFR αD-βE loop, while LOV2 insertions in this region location did not initiate allostery as quantified either *in vitro* or *in vivo* [20, 21]. This suggests different structural requirements for establishing and tuning allostery in this system (and possibly others): here allostery seems to be more easily introduced at evolutionarily conserved and co-evolving sites, but once established, can be optimized through less conserved sector-peripheral residues.

Though our work focuses on a synthetic allosteric fusion, our results are broadly consistent with an emerging body of work characterizing allostery-influencing mutations in natural proteins. Together, these data point to a model in which mutations at evolutionarily conserved positions exert large (and often disruptive) effects on function while allostery is tuned at less conserved surface sites. For example, Leander et al. recently used deep mutational scanning to map the pattern of compensatory mutations that rescued allosteric function for non-allosteric tetracycline repressor (TetR) variants [56]. In that study a “disrupt-and-restore” strategy was used: an already-allosteric system was inactivated and deep mutational scanning was then used to identify compensatory mutations. While there are significant differences between rescuing a deficient variant and the optimization of a novel allosteric construct, they likewise found that the mutations at highly conserved sites were often disruptive to stability and function, while allostery-rescuing mutations occurred at weakly conserved and structurally distributed sites [56]. Similarly, mutations at “rheostat” sites – weakly conserved positions distal to the site of regulation – were found to modulate allosteric control in human liver pyruvate kinase and the lactose repressor protein (lacI) [57, 58]. Intriguingly, the association of allostery enhancing mutations with the protein surface hints at a possible role for solvent – and more specifically the protein hydration layer – in tuning regulation.

The finding that the allostery initiated upon naïve fusion of the DHFR and LOV2 domains can be further enhanced by single mutations implies a path to improved allosteric dynamic range by stepwise mutagenesis and selection. Three of the most allostery enhancing mutations could be combined to yield a near-additive improvement in regulatory dynamic range. This has interesting implications for both evolved and engineered allosteric systems. Given that standing mutational variation in a population is more likely at weakly conserved surface sites (particularly under less stringent selection conditions), this could provide a means for generating variation in allosteric regulation upon a domain fusion event. Moreover, while engineering studies sometimes use mutations near the domain insertion site to optimize regulation, our results suggest that diffuse surface site mutations would present a more effective alternative. Whether by engineering or evolution, it seems that mutations at weakly conserved and structurally distributed residues can provide a path to the optimization of regulation.

## MATERIALS AND METHODS

### EXPERIMENTAL MODEL AND SUBJECT DETAILS

#### Escherichia coli expression and selection strains

ER2566 Δ*folA* Δ*thyA E. coli* were used for all growth *in vivo* growth rate measurements; this strain was a kind gift from Dr. Steven Benkovic and is the same used in Reynolds et al., 2011 and Thompson et al., 2020 [21, 27]. XL1-Blue *E. coli* (genotype: *recA1 endA1 gyrA96 thi-1 hsdR17 supE44 relA1 lac* [F’ *proAB lacI^q^ZΔM15* Tn*10*(Tet^r^)]) from Agilent Technologies were used for cloning, mutagenesis, and plasmid propagation. BL21(DE3) *E. coli* (genotype: *fhuA2 [lon] ompT gal* (*λ DE3*) *[dcm] ΔhsdS. λ DE3 = λ sBamHIo ΔEcoRI-B int::(lacI::PlacUV5::T7 gene1) i21 Δnin5*) from New England Biolabs were used for protein expression.

### METHOD DETAILS

#### DHFR Saturation Mutagenesis Library Construction

The construction of the DHFR-LOV2 saturation mutagenesis library was done as described in Thompson et al., 2020 [27]. Four sublibraries were generated to cover the entire mutational space of *E. coli* DHFR: positions 1–40 (sublibrary1, SL1), positions 41–80 (sublibrary2, SL2), positions 81–120 (sublibrary3, SL3), and positions 121–159 (sublibrary4, SL4) Inverse PCR with NNS mutagenic primers (N = A/T/G/C, S = G/C) was done at every position in DHFR to produce all amino acid substitution. The vector with DHFR-LOV2 121 and TYMS in a pACYC-Duet vector was described in Reynolds et al., 2011 [21].

The NNS primers were phosphorylated with T4 polynucleotide kinase (NEB, cat#M0201S). 20 μL phosphorylations was prepared according to the following recipe: 16.5 μL sterile water, 2 μL T4 ligase buffer, 0.5 μL T4 PNK enzyme, and 1 μL 100 μM NNS primers. The reactions were then heated at 37°C for 1 hour and 65°C for 20 minutes.

PCR reactions were set up using 2x Q5 mastermix (NEB, cat#M0492), 10 ng of plasmid template, and 500 nM forward and reverse primers. PCR was performed in the following steps: 1) 98°C for 30 seconds, 2) 98°C for 10 seconds, 3) 55°C for 30 seconds, 4) 72°C for 2 minutes, 5) return to step 2 for 22 cycles, 6) 72°C for 5 minutes. 25 μL of PCR reaction was mixed with 1 μL of DpnI (NEB, cat#R0176) at 37°C for 4 hours. The samples were then purified by gel extraction and a DNA Clean & Concentrator −5 kit (Zymo Research, cat#D4014). PCR product solution were then phosphorylated with a second round of T4 PNK: 100 μL of gel-extracted PCR product,12 μL of 10x T4 ligase buffer, 5 μL of T4 PNK, 5 μL of sterile water and were incubated at 37°C for 1 hour with 90°C for 30 seconds. The reactions were ligated with 100 μL PNK phosphorylated PCR product, 15 μL T4 ligase (NEB, cat#M0202S), 30 μL T4 ligase buffer and, 155 μL sterile water. The reaction was incubated at room temperature for 24 hours.

The concentration of each reaction was quantified by gel densitometry (ImageJ) and combined in equimolar ratios to form sublibraries. The library was divided up into four sublibraries with sublibrary 1 covering positions 1-40, sublibrary 2 covering positions 41-80, sublibrary 3 covering positions 81-120, and sublibrary 4 covering positions 121-150. Sublibraries were transformed into electrocompetent XL1-Blue *E. Coli* using a MicroPulser Electroporator (Bio Rad) and gene pulser cuvettes (Bio Rad, cat#165-2089). Cultures were miniprepped using a GeneJET plasmid miniprep kit (Thermo Scientific, cat#K05053). Library completeness was verified by deep sequencing on a MiSeq (Illumina).

#### Growth Rate Measurements in the Turbidostat for DHFR DL121 Mutant Library

DHFR DL121 sublibraries were transformed into ER2566 Δ*folA* Δ*thyA E. coli* by electroporation using a MicroPulser Electroporator (Bio Rad) and gene pulser cuvettes (Bio Rad, cat#165-2089). Cultures were grown overnight at 37°C in GM9 minimal media (93.0 mM Sodium (Na^+^), 22.1 mM Potassium (K^+^), 18.7 mM Ammonium (NH_4_), 1.0 mM Calcium (Ca^2+^), 0.1 mM Magnesium (Mg^2+^), 29.2 mM Chloride (Cl^-^), 0.1 mM Sulfate (SO_4_^2-^), and 42.2 mM Phosphate (PO_4_^3-^), 0.4% glucose) pH 6.50, containing 50 μg/mL thymidine and 30 μg/mL chloramphenicol (Sigma, cat#C0378-5G) as well as folA mix which contains 38 μg/mL glycine (Sigma, cat#50046), 75.5 μg/mL L-methionine (Sigma, cat#M9625) 1 μg/mL calcium pantothenate (Sigma, cat#C8731), and 20 μg/mL adenosine (Sigma, cat#A9251). Four hours before the start of the experiment the overnight culture was diluted to an optical density of 0.1 at 600nm in GM9 minimal media containing 50 μg/mL thymidine and 30 μg/mL chloramphenicol and incubated for four hours at 30°C. The cultures were centrifuged at 2,000RCF for 10 minutes and resuspended in the experimental conditions of GM9 minimal media containing 1 μg/mL thymidine and 30 μg/mL chloramphenicol. This was repeated two more times. The cultures were then back-diluted to an OD600 of 0.1 in 16 mL/vial of media. The turbidostat described in Toprak et al., 2013 was used in continuous culture (turbidostat) mode with a clamp OD600 of 0.15 and a temperature of 30°C. Each vial had a stir bar. Vials designated as “lit” had one 5V blue LED active. The optical density was continuously monitored throughout the experiment. 1mL samples were taken at the beginning of selection (0 hour) and at 4, 8, 12, 16, 20, and 24 hours into selection and were centrifuged at 21,130 RCF for 5 minutes at room temperature with the pellet being stored at −20°C for sequencing sample preparation.

#### Growth Rate Measurements in the Turbidostat for DHFR Control Library

Wild-type DHFR, 12 DHFR point mutants (D27N, F31V, F31Y, F31Y-L54I, G121V, G121V-F31Y, G121V-M42F, L54I, L54I-G121V, M42F, and W22H), and three chimeric DHFR-LOV2 fusion constructs (DL116, DL121, and DL121-C450S) each in a pACYC-Duet vector with TYMS as described in Reynolds et al., 2011 were transformed into ER2566 Δ*folA* Δ*thyA E. coli* by electroporation using a MicroPulser Electroporator (Bio Rad) and gene pulser cuvettes (Bio Rad, cat#165-2089) [21]. Cultures were grown overnight at 37°C in GM9 minimal media (93.0 mM Sodium (Na^+^), 22.1 mM Potassium (K^+^), 18.7 mM Ammonium (NH_4_), 1.0 mM Calcium (Ca^2+^), 0.1 mM Magnesium (Mg^2+^), 29.2 mM Chloride (Cl^-^), 0.1 mM Sulfate (SO_4_^2-^), and 42.2 mM Phosphate (PO_4_^3-^), 0.4% glucose) pH 6.50, containing 50 μg/mL thymidine and 30 μg/mL chloramphenicol (Sigma, cat#C0378-5G) as well as folA mix which contains 38 μg/mL glycine (Sigma, cat#50046), 75.5 μg/mL L-methionine (Sigma, cat#M9625) 1 μg/mL calcium pantothenate (Sigma, cat#C8731), and 20 μg/mL adenosine (Sigma, cat#A9251). Four hours before the start of the experiment the overnight culture was diluted to an optical density of 0.1 at 600nm in GM9 minimal media containing 50 μg/mL thymidine and 30 μg/mL chloramphenicol and incubated for four hours at 30°C. The cultures were centrifuged at 2,000RCF for 10 minutes and resuspended in the experimental conditions of GM9 minimal media containing 1 μg/mL thymidine and 30 μg/mL chloramphenicol. This was repeated two more times. The cultures were then back-diluted to an OD600 of 0.1 and pooled at equal (1/16^th^) ratios and aliquoted into four “dark” and four “lit” vials with 16ml culture. The turbidostat described in Toprak et al., 2013 was used in continuous culture (turbidostat) mode with a clamp OD600 of 0.15 and a temperature of 30°C. Each vial had a stir bar. Vials designated as “lit” had one 5V blue LED active. The optical density was continuously monitored throughout the experiment. 1mL samples were taken at the beginning of selection (0 hour) and at 4, 8, 12, 16, 20, and 24 hours into selection and were centrifuged at 21,130 RCF for 5 minutes at room temperature with the pellet being stored at −20°C for sequencing sample preparation.

#### Plate reader assay for E. coli growth

Single point mutant DHFR-D27N, DL121 chimeric protein, and DL121 with a point mutant D27N each in a pACYC-Duet vector with TYMS as described in Reynolds et al., 2011 were transformed into ER2566 Δ*folA* Δ*thyA E. coli* by electroporation using a MicroPulser Electroporator (Bio Rad) and gene pulser cuvettes (Bio Rad, cat#165-2089) [21]. Cultures were grown overnight at 37°C in GM9 minimal media (93.0 mM Sodium (Na^+^), 22.1 mM Potassium (K^+^), 18.7 mM Ammonium (NH_4_), 1.0 mM Calcium (Ca^2+^), 0.1 mM Magnesium (Mg^2+^), 29.2 mM Chloride (Cl^-^), 0.1 mM Sulfate (SO_4_^2-^), and 42.2 mM Phosphate (PO_4_^3-^), 0.4% glucose) pH 6.50, containing 50 μg/mL thymidine and 30 μg/mL chloramphenicol (Sigma, cat#C0378-5G) as well as folA mix which contains 38 μg/mL glycine (Sigma, cat#50046), 75.5 μg/mL L-methionine (Sigma, cat#M9625) 1 μg/mL calcium pantothenate (Sigma, cat#C8731), and 20 μg/mL adenosine (Sigma, cat#A9251). Four hours before the start of the experiment the overnight culture was diluted to an optical density of 0.1 at 600nm in GM9 minimal media containing 50 μg/mL thymidine and 30 μg/mL chloramphenicol and incubated for four hours at 30°C. The cultures were centrifuged at 2,000RCF for 10 minutes and resuspended in the experimental conditions of GM9 minimal media containing either 0, 1, or 50 μg/mL thymidine and 30 μg/mL chloramphenicol. The cells were centrifuged and resuspended two more times. The cultures were then back-diluted to an OD600 of 0.005 into 96-well plates with 6 replicates each.

#### Next Generation Sequencing Amplicon Sample Preparation

Cell pellets were lysed by the addition of 10 μL sterile water, mixed by pipetting, and incubated at 98°C for 5 minutes. 1 μL of this was then combined with 5 μL Q5 buffer (NEB, cat#M0491 S), 0.5 μL 10mM DNTP (Thermo Scientific, cat#R0192), 2.5 μL of 10mM forward and reverse primers specific to the sublibrary and containing the TruSeq adapter sequence (Appendix 1: SL1V2, SL2V2, SL3V2, SL4V2, DL121CLV3F, and DL_WTTS_R3), 0.25 μL of Q5 enzyme (NEB, cat#M0491S) and 13.25 μL of sterile water. These samples were then heated at 98°C for 90 seconds and then cycled through 98°C for 10 seconds 63-65°C (sublibrary 1: 66°C, sublibrary 2: 63°C, sublibrary 3: 64°C, and sublibrary 4: 65 °C) for 15 seconds and then 72°C for 15 seconds, repeating 20 times with a final 72°C heating for 120 seconds in a Veriti 96 well thermocycler (Applied Biosystems). These samples were then amplified using TruSeq PCR reactions with a unique combination of i5/i7 indexing primers for each timepoint. 1 μL of this PCR reaction was then combined with 5 μL Q5 buffer (NEB, cat#M0491S), 0.5 μL 10mM DNTP (Thermo Scientific, cat#R0192), 2.5 μL of 10mM forward and reverse primers, 0.25 μL of Q5 enzyme (NEB, cat#M0491S) and 13.25 μL of sterile water. These samples were then heated at 98°C for 30 seconds and then cycled through 98°C for 10 seconds 55°C for 10 seconds and then 72°C for 15 seconds, repeating 20 times with a final 72°C heating for 60 seconds in a Veriti 96 well thermocycler (Applied Biosystems). Amplified DNA from i5/i7 PCR reaction was quantified using the picogreen assay (Thermo Scientific, cat#P7589) on a Victor X3 multimode plate reader (Perkin Elmer) and the samples were mixed in an equimolar ratio. The DNA was then purified by gel extraction and a DNA Clean & Concentrator −5 kit (Zymo Research, cat#D4014). DNA quality was determined by 260nm/230nm and 260nm/280nm ratios on a DS-11+ spectrophotometer (DeNovix) and concentration was determined using the Qubit 3 (Thermo Scientific). Pooled samples were sent to GeneWiz where they were analyzed by TapeStation (Agilent Technologies) and sequenced on a HiSeq 4000 sequencer (Illumina) with 2×150bp dual index run with 30% PhiX spike-in yielding 1.13 billion reads. The control library was sequenced in-house using a MiSeq sequencer (Illumina) with 2×150bp dual index 300 cycle MiSeq Nano Kit V2 (Illumina cat#15036522) with 20% PhiX (Illumina cat#FC-110-3001) spike-in yielding 903,488 reads.

#### DHFR Chimeric Expression Constructs

The *E. coli* DHFR LOV2 fusion was cloned as an NcoI/XhoI fragment into the expression vector pHIS8-3 [20, 21]. Point mutants were engineered into the DHFR gene using QuikChange II site-directed mutagenesis kits (Agilent cat#200523) using primers specified in Appendix 1. All DHFR/LOV2 fusions for purification were expressed under control of a T7 promoter, with an N-terminal 8X His-tag for nickel affinity purification. The existing thrombin cleavage site (LVPRGS) following the His-tag in pHIS8-3 was changed to a TEV cleavage site using restriction-free PCR to improve the specificity of tag removal [59]. All constructs were verified by Sanger DNA sequencing.

#### Protein expression and purification

DHFR-LOV2 chimeric proteins were expressed in BL21(DE3) *E. coli* grown at 30°C in Terrific Broth (12g/L Tryptone, 24g/L yeast extract, 4mL/L glycerol, 17mM KH_2_PO_4_, and 72mM K_2_HPO_4_). Protein expression was induced when the cells reached an absorbance at 600nm of 0.7 with 0.25 mM IPTG, and cells were grown at 18°C overnight. Cell pellets were lysed by sonication in binding buffer (500 mM NaCl, 10 mM imidazole, 50 mM Tris-HCL, pH 8.0) added at a volume of 5ml/g cell pellet. Next the lysate was clarified by centrifugation and the soluble fraction was incubated with equilibrated Ni-NTA resin (Qiagen cat#4561) for 1 hour at 4°C. After washing with one column volume of wash buffer (300 mM NaCl, 20mM imidazole, 50 mM Tris-HCL, pH 8.0) the DHFR-LOV2 protein was eluted with elution buffer (1M NaCl, 250 mM imidazole, 50mM Tris-HCL, pH 8.0) at 4°C. Eluted protein was dialyzed into dialysis buffer (300 mM NaCl, 1% glycerol, 50 mM Tris-HCl, pH 8.0) at 4°C overnight in 10,000 MWCO Thermo protein Slide A Lyzer (Fisher Scientific cat#PI87730). Following dialysis, the protein was then purified by size exclusion chromatography (HiLoad 16/600 Superdex 75 pg column, GE Life Sciences cat#28989333). Purified protein was concentrated using Amicon Ulta 10k M.W. cutoff concentrator (Sigma cat#UFC801024) and flash frozen using liquid N2 prior to enzymatic assays.

#### Steady state Michaelis Menten Measurements

The protein was spun down at 21,130 RCF at 4°C for 10 minutes and the supernatant was moved to a new tube with any pellet being discarded. The concentration of the protein was quantitated by A280 using a DS-11 + spectrophotometer (DeNovix) with an extinction coefficient of 44920 mM^-1^ cm^-1^. The parameters k_cat_ and K_m_ under Michaelis-Menten conditions were determined by measuring the initial velocity for the depletion of NADPH as measured in absorbance at 340nm, with an extinction coefficient of 13.2 mM^-1^ cm^-1^. This is done in a range of substrate concentrations with a minimum of 8 data points around 4 K_m_, 2 K_m_, 1.5 K_m_, K_m_, 0.8 K_m_, 0.5 K_m_, 0.25 K_m_ and 0. The initial velocities (slope of the first 15 seconds) were plotted vs. the concentration of Dihydrofolate and fit to a Michaelis Menten model using non-linear regression in GraphPad Prism 7. The reactions are run in MTEN buffer (50mM 2-(N-morpholino)ethanesulfonic acid, 25mM tris base, 25mM ethanolamine, 100mM NaCl) pH 7.00, 5mM Dithiothreitol, 90μM NADPH (Sigma-Aldrich cat#N7505) quantitated by A340. Dihydrofolate (Sigma-Aldrich cat#D7006) is suspended in MTEN buffer pH 7.00 with 0.35% β-mercaptoethanol and quantitated by A282 with an extinction coefficient of 28 mM^-1^ cm^-1^. Depletion of NADPH is observed in 1 mL cuvettes with a path length of 1cm in a Lambda 650 UV/VIS spectrometer (Perkin Elmer) with attached water Peltier system set to 17°C. Lit samples are illuminated for at least 2 minutes by full spectrum 125 watt 6400K compact fluorescent bulb (Hydrofarm Inc. cat#FLC125D). Dark samples were also exposed to the light in the same way as the lit samples but were in opaque tubs.

#### Spectrophotometry of the LOV2 chromophore

The spectra of the LOV2 chromophore is determined with a Lambda 650 UV/VIS spectrometer (Perkin Elmer) at 350-550nm using paired 100 μL Hellma ultra micro cuvettes (Sigma cat#Z600350-1EA) with a path length of 1cm. Purified protein in was diluted (when possible) to 20 μM in MTEN buffer pH 7.00 with 0.35% β-mercaptoethanol The lit samples are illuminated for at least 2 minutes by full spectrum 125 watt 6400K compact fluorescent bulb (hydrofarm Inc). Relaxation of the lit state chromophore is observed in the Lambda 650 UV/VIS spectrometer (Perkin Elmer) at 447nm (dark peak) using paired 100 μL Hellma ultra micro cuvettes (Sigma cat#Z600350-1EA) with a path length of 1cm.

### QUANTIFICATION AND STATISTICAL ANALYSIS

#### Next Generation Sequencing

The sequencing data analysis can be divided into two portions: 1) Read Joining, Filtering and Counting, followed by 2) Calculating Relative Fitness and Final Filtering. We describe each step below; all code was implemented in Bash shell scripting or Python 3.6.4

#### Read Joining, Filtering, and Counting

The data analysis began with unjoined illumina fastq.gz files separated by index (generated by GeneWiz). The forward and reverse reads were combined using usearch v11.0.667 using the i86linux32 package. The commands given to usearch are contained in the script **UCOMBINER.bsh**.

Reads of each paired fastq file are identified and quality checked using the script **DL121_fastq_analysis.py**. Mutant nucleotide counts and number of wild type reads are stored in a dictionary where the read count is separated by file name (vial and timepoint eg: T2V3) and sublibrary. If any nucleotide in the coding region is below a qscore cutoff of 30, that read is discarded. Counts of every nucleotide are saved in a text file by timepoint and vial.

Converting nucleotide variation to amino acid count as well as probabilistic sequencer error correction is done by the **Hamming_analysis.ipynb** script. Given the probabilistic nature of base calling on the Illumina platform, one can expect a number of reads that were errantly called. For each codon the expected number of reads due to sequencing noise was calculated with the formula:

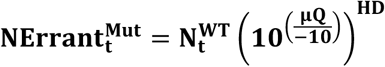

The number of errant mutants 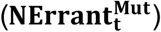 **can** be calculated from the number of observed wild type 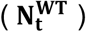, the average Q score of the sequencing run **μQ**, and the hamming distance (**HD**) or number of mutations away from. The number of errant mutants then subtracted from the actual mutant count. In addition to the number of observed wild type, this is calculated for every possible mutation observed, up to the 31 other nucleotide codons, (NNK codons are discarded due to the nature of library construction). Once the total number of errant reads are calculated and subtracted from the mutant and wild type counts, they are then converted into the amino acid sequence and are saved into text files.

These files are then used to load information for calculation of growth rate and allostery:

#### Calculating Relative Fitness and Final Filtering

**Growth_Rate_and_Allostery.ipynb** was the python script used for this analysis. Relative frequency was calculated as follows:

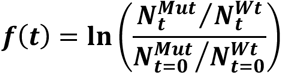

Variant frequencies 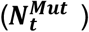 were determined relative to WT 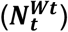 and normalized to the initial frequency distribution at t=0. The relative growth rate then calculated by linear regression of these normalized frequencies. Light dependence was calculated as the difference between lit vs. dark growth rates. Variant frequency was only calculated if there were more than 50 mutant reads at time zero. Definitions for sector identity, conservation values, and surface identity used in **SectorSurfaceDefinitions.ipynb** are the same as those from *Reynolds 2011* [21]. Accessible surface area was calculated using MSMS, using a probe size of 1.4Å and excluding water as well as heteroatoms [60]. Values for total surface areas were taken from *Chothia 1975* [61]. Together these were used to calculate relative solvent accessible surface area, and 25% was used as a cutoff for “surface”. A surface site is considered to contact the sector if the atoms comprising the peptide bond contact *any* sector atoms. Contact is defined as the sum of the atom’s Pauling radii + 20%.

To determine significant allosteric mutations, a p-value for each mutation was computed by unequal variance t-test under the null hypothesis that the lit and dark replicate measurements have equal means. Two cutoffs were used, a standard cutoff of P=0.05, and a more stringent cutoff that is adjusted to consider multiple hypothesis testing. A multiple-hypothesis testing adjusted p-value of P=0.016 was determined by Sequential Goodness of Fit [49]. General analysis and figures made from this data are performed in **allostery_analysis.ipynb.**

## Appendix 1 – Primer table

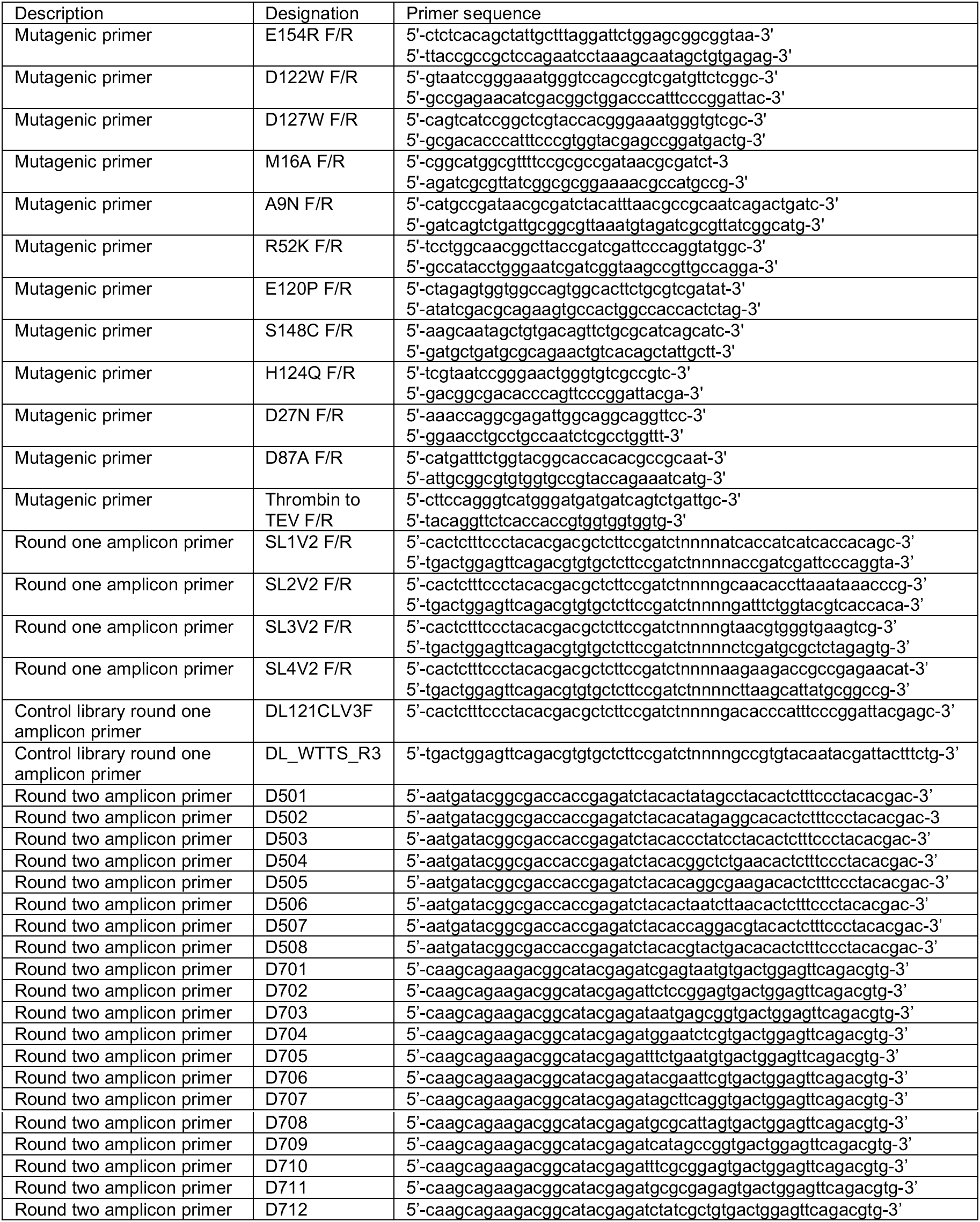

## ACKNOWLEDGEMENTS

The authors are grateful to Dr. Tanja Kortemme for facilitating our collaboration with Samuel Thompson. We also acknowledge Dr. Elliott Ross and Dr. Rama Ranganathan for thoughtful discussion and feedback. We thank Christine Ingle for her assistance with DHFR purification and kinetics protocols, and other members of the Reynolds lab for comments on the manuscript and discussions throughout the development of this work.

## FUNDING

This work was supported by NSF Grant # 1942354 to KAR, and in part by the Gordon and Betty Moore Foundation’s Data Driven Discovery Initiative through grant GBMF4557 to KAR.

## AUTHOR CONTRIBUTIONS

JWM and KAR conceptualized the work and designed experiments. JWM performed the majority of experiments with assistance from MAXR (expression and purification of DL121 point mutants), AB (collection of the growth assay “calibration curve” data in Figure 2) and SMT (construction of the deep mutational scanning library). JWM and KAR performed all data analysis. JWM and KAR wrote the original draft. MAXR, AB, and SMT contributed to manuscript review and editing. KAR provided supervision and funding acquisition.

## COMPETING INTERESTS

The authors have no competing interests to declare.

## DATA AND MATERIALS AVAILABILITY

Plasmid constructs for DL121 will be deposited with addgene during the manuscript review process.

Code for analysis of the deep mutational scanning data is available through Github: https://github.com/reynoldsk/allostery-in-dhfr

The sequencing data is available in FASTQ format: https://www.ncbi.nlm.nih.gov/bioproject/PRJNA706683 https://trace.ncbi.nlm.nih.gov/Traces/sra/?study=SRP309306

## Notes

https://github.com/reynoldsk/allostery-in-dhfr

https://www.ncbi.nlm.nih.gov/bioproject/PRJNA706683

### Competing Interest Statement

The authors have declared no competing interest.

