## Supplemental Information for "Structurally Distributed Surface Sites Tune Allosteric Regulation"

**Contents**:

### I. Supplementary Figures

Figure S1: Deep mutational scanning library completeness – heatmap of sequencing counts for all mutants

Figure S2: Deep mutational scanning library completeness – distribution of sequencing counts for all mutants

Figure S3: Reproducibility across biological replicates

Figure S4: Heatmaps of relative growth rates under dark and lit conditions; heatmap of calculated allostery

Figure S5: Growth rate measurements for DL121-D27N

Figure S6: Relationship between catalytically inactivating mutations and evolutionarily conserved positions

Figure S7: Distribution of mutational effects on allosteric regulation.

Figure S8: Steady state kinetics measurements for select mutants in the light and dark.

Figure S9: Spectroscopic characterization of LOV2 activation for select DL121 mutants.

Figure S10: Relaxation rate of the LOV2 chromophore for select DL121 mutants.

Figure S11: Correlation between *in vivo* allostery and *in vitro* steady state kinetics parameters

Figure S12: Characterization of the DL121- M16A,H124Q and DL121- M16A, D87A, H124Q mutants

### II. Supplementary Tables

Table S1. Steady-state kinetic parameters for select DL121 mutants

Table S2. Statistical association of catalytically inactivating mutations and the sector

Table S3: Statistical association of catalytically inactivating mutations and conservation

Table S4. Statistical association of allosteric mutations and the sector

Table S5. Statistical association of allosteric mutations and the protein surface

Table S6: Statistical association of allosteric mutations and sector connected (including positions within sector) surface sites.

Table S7: Statistical association of allosteric mutations and sector connected (excluding positions within sector) surface sites.

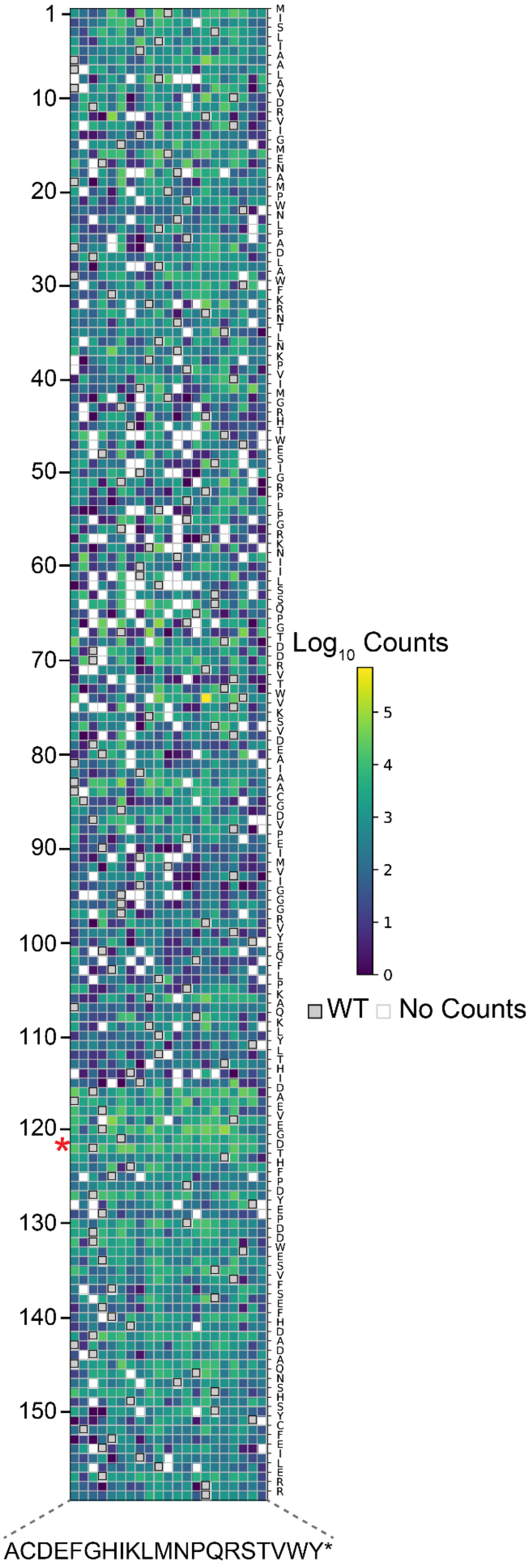


**Figure S1:** Deep mutational scanning library completeness – heatmap of counts for all mutants. $Log_{10}$(counts) of all possible mutations in DHFR domain of DL121 chimeric protein library at time point zero. The y axis corresponds to positions on *E. coli* DHFR domain as numbered in PDB ID: 1RX2. A red star indicates the location of the LOV2 domain insertion. The x axis corresponds to possible mutations. Wild type residues are shown in grey; positions with no counts are shown in white.

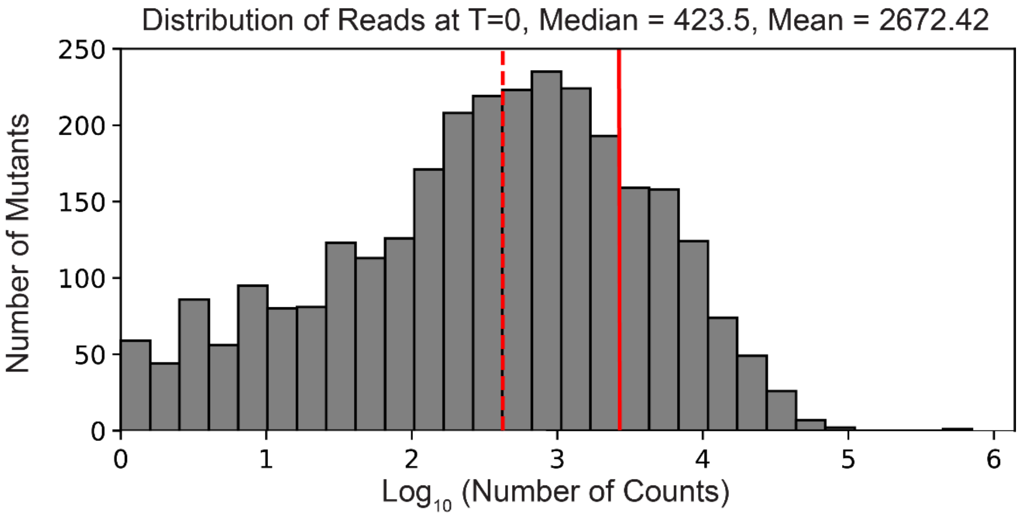


**Figure S2:** Deep mutational scanning library completeness – distribution of counts for all mutants. A histogram of the number of counts per mutant at time point zero. The median and mean number of counts is shown as a dashed and solid red line, respectively.

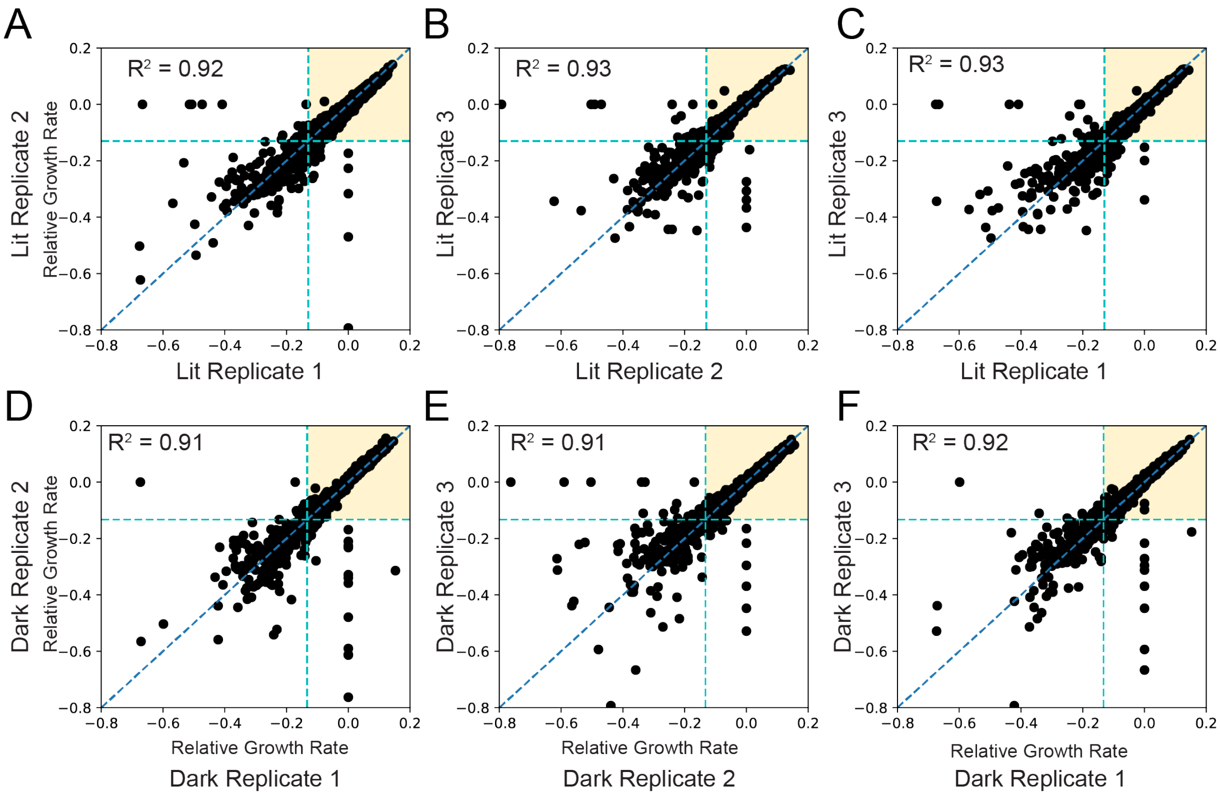


**Figure S3:** Reproducibility across biological replicates. The relative growth rate (see methods) for each mutant is compared across all three lit (A-C) and dark (D-F) replicates. The line of best fit is indicated with a blue dashed line. The teal dashed lines represent the growth rate of DL121-D27N; mutants with a relative growth rate below this cutoff were considered near catalytically inactive and excluded from analysis of allostery.

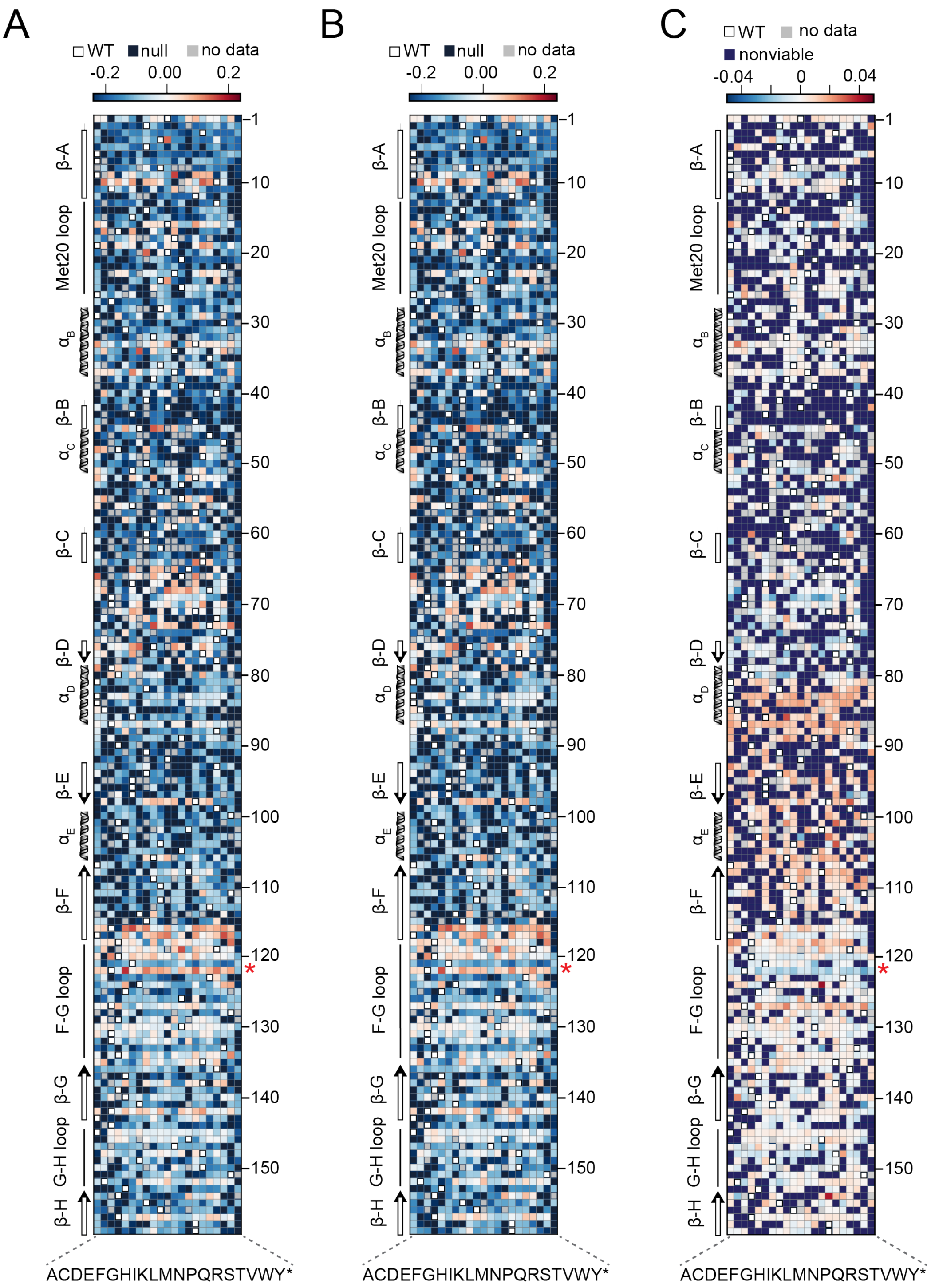


**Figure S4:** Heatmaps of relative growth rates and allostery. A) Relative growth rate in the dark. B) Relative growth rate in the light. C) Allostery. In panels A-B, blue and red indicate mutations with deleterious and beneficial effects on growth rate respectively. In panel C, blue indicates allostery disrupting mutations, and red indicates allostery enhancing mutations. In all three panels, white squares with black outlines mark the WT residue at each position. Mutations missing from the library ('no data') are colored grey, and mutations that did not have sufficient counts for at least three time points ('null data', no relative growth rate could be fit) are colored navy.

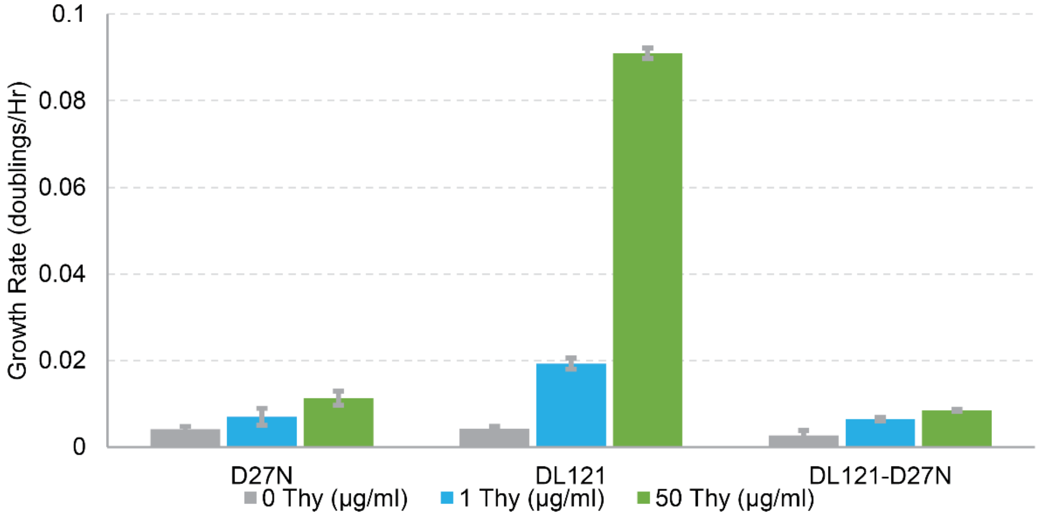


**Figure S5:** Growth rate measurements for DL121-D27N. Comparison of growth rates as doublings per hour for three enzymes: nonchimeric *E. coli* DHFR with a D27N mutation (rendering it catalytically inactive), the unmutated fusion protein, DL121, and DL121 combined with the D27N mutation. All three mutants were grown in a 96-well plate in M9 media supplemented with either no thymidine, 1µg/ml thymidine (the same media conditions as the experiments in this work), or 50 µg/ml thymidine at 30°C. Error bars represent standard deviation across six replicates.

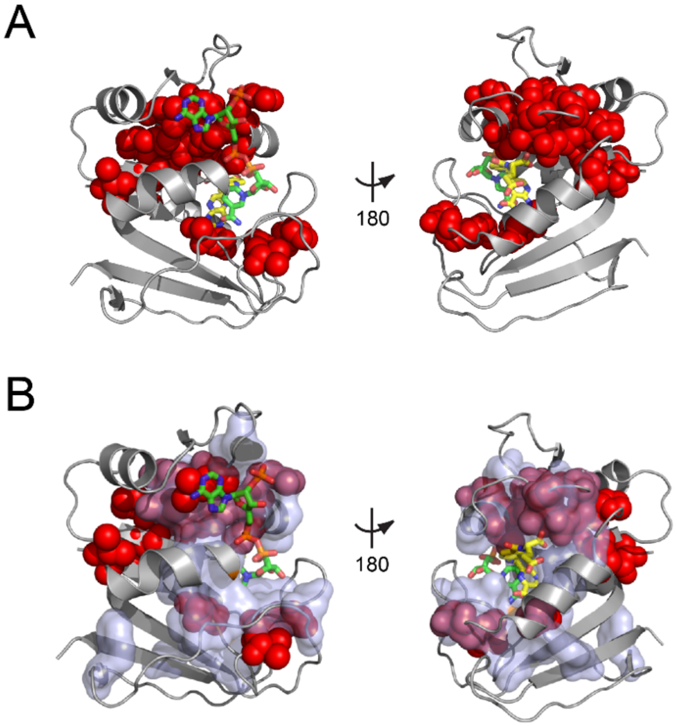


C

Conserved Positions

| Low Growth | Y | N | |
|---|---|---|---|
| Y | 413 | 834 | 1247 |
| N | 280 | 1268 | 1548 |
| | 693 | 2102 | 2795 |

$P = 8.7 \times 10^{-20}$

**Figure S6:** Relationship between catalytically inactivating mutations and evolutionarily conserved positions. A) Structural distribution of positions enriched for mutations with growth rates as low as or lower than DL121 D27N (indicated with red spheres). The DHFR backbone is in grey cartoon, the folate substrate in yellow sticks, and the NADP co-factor in green sticks. B) Relationship of evolutionarily conserved positions (light blue surface) to positions enriched for growth-rate disrupting mutations (red spheres, same as in A). C) A contingency table summarizes the overlap between conserved positions, and the mutations that yield low growth (growth rate <= DL121 D27N).

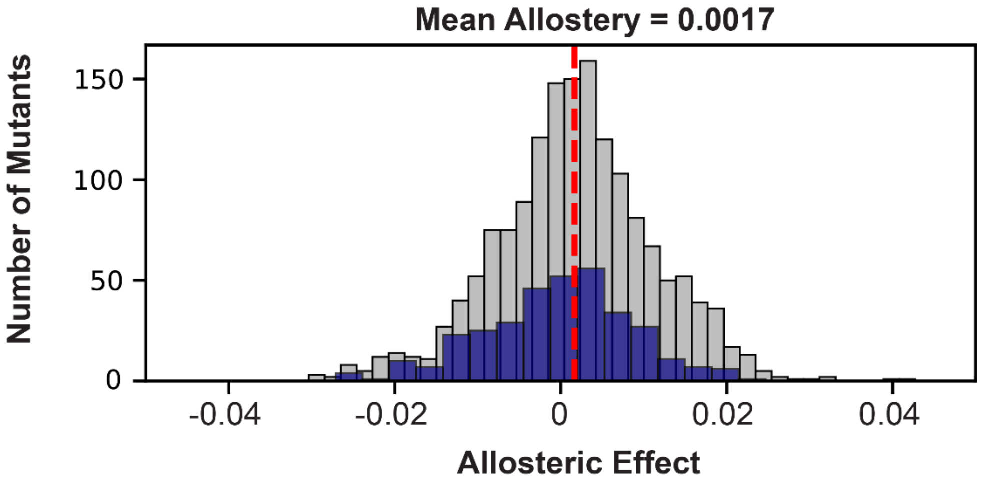


**Figure S7**. Distribution of mutational effects on allosteric regulation. The allosteric effect of all viable mutants is shown in grey with the mean allosteric effect of 0.0017 shown as a red dotted line. The allosteric effect of viable mutants in the sector is shown overlaid in blue. The mean allosteric effect of sector positions is -0.0005. The cutoff for sector identity used is a p-value of 0.01 as calculated in *Reynolds 2011* [21].

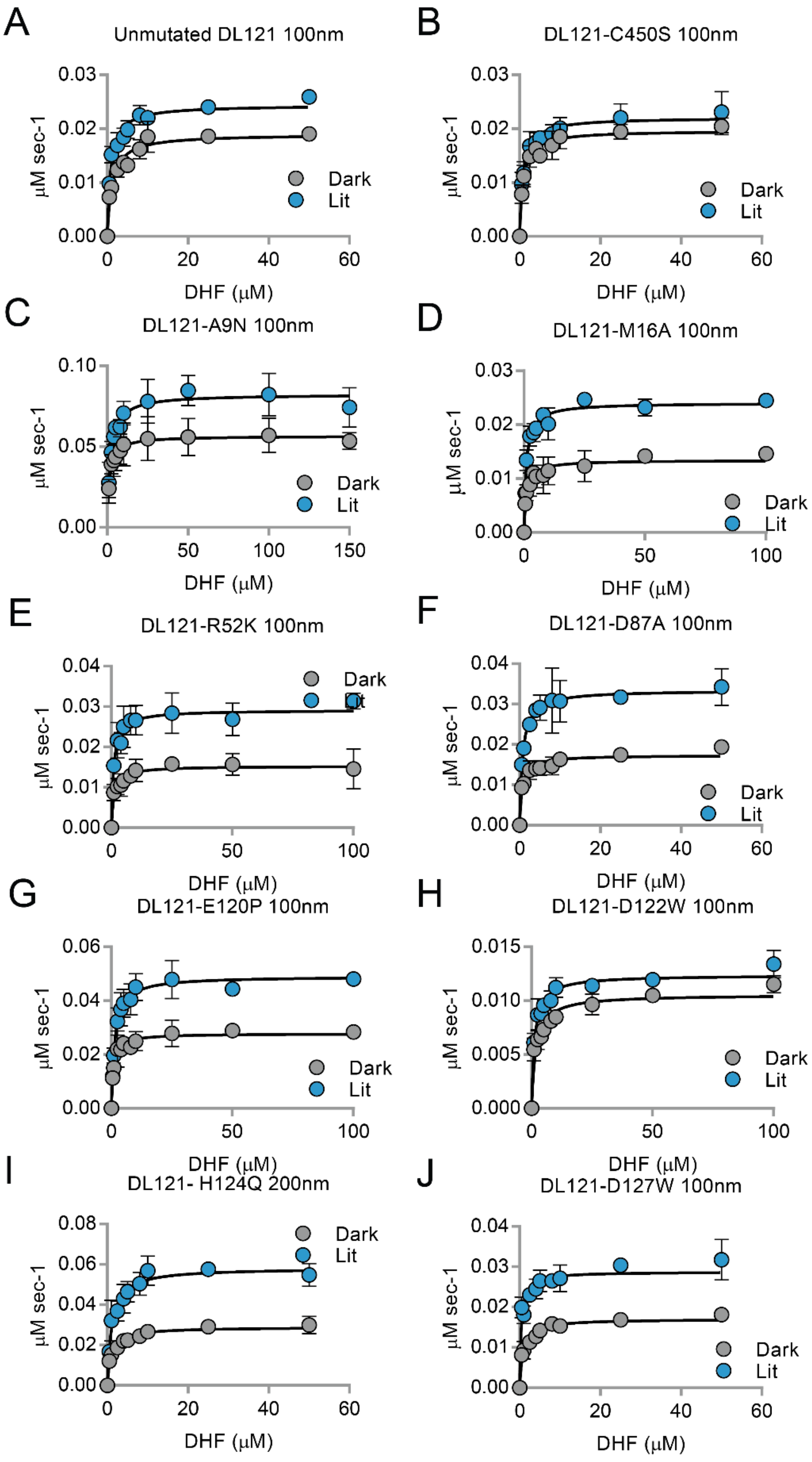
A
Unmutated DL121 100nm
B
DL121-C450S 100nm
C
DL121-A9N 100nm
D
DL121-M16A 100nm
E
DL121-R52K 100nm
F
DL121-D87A 100nm
G
DL121-E120P 100nm
H
DL121-D122W 100nm
I
DL121- H124Q 200nm
J
DL121-D127W 100nm
μM sec-1
DHF (μM)
Dark
Lit

**Figure S8 (previous page):** Steady state kinetics measurements for select mutants in the light and dark. Initial velocity vs. substrate (dihydrofolate) concentration for the purified DL121 chimeric protein, allosterically inactivated DL121-C450S and 9 point mutations to the DHFR domain of DL121. Lit (blue) and dark (grey) conditions are shown with error bars representing standard deviation across three replicates. The $k_{cat}$, $K_M$, catalytic efficiency and associated error are reported in Table S1.

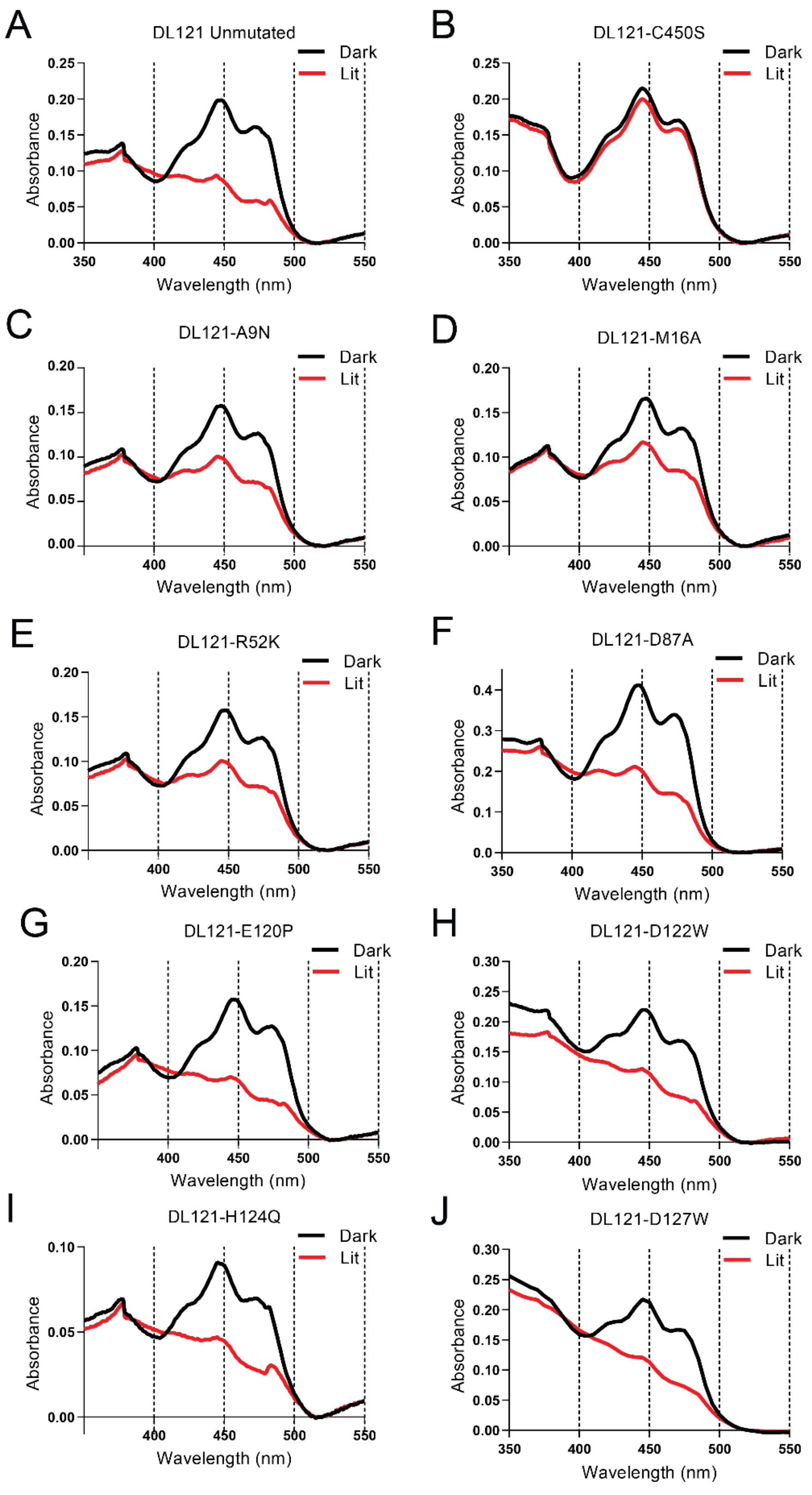
A
DL121 Unmutated
B
DL121-C450S
C
DL121-A9N
D
DL121-M16A
E
DL121-R52K
F
DL121-D87A
G
DL121-E120P
H
DL121-D122W
I
DL121-H124Q
J
DL121-D127W
Dark
Lit
Absorbance
Wavelength (nm)

**Figure S9 (previous page):** Spectroscopic characterization of LOV2 activation for select DL121 mutants. The absorbance of purified DL121 chimeric protein, allosterically inactivated DL121-C450S and 9 point mutations to the DHFR domain of DL121. Lit state absorbance spectra (red line) were measured after illumination for at least 2 minutes by full spectrum 125 watt 6400K fluorescent lamp (Hydrofarm Inc). Dark conditions are taken under the same conditions but using opaque tubes when the sample was placed under the lamp. With the exception of the DL121-C450S mutant, all show a characteristic spectral shift upon light stimulation consistent with an active LOV2 domain. Formation of a covalent FMN-thiol adduct in the LOV2 domain upon light exposure causes the 447nm peak in the dark state to shift to 390 nm in the light.

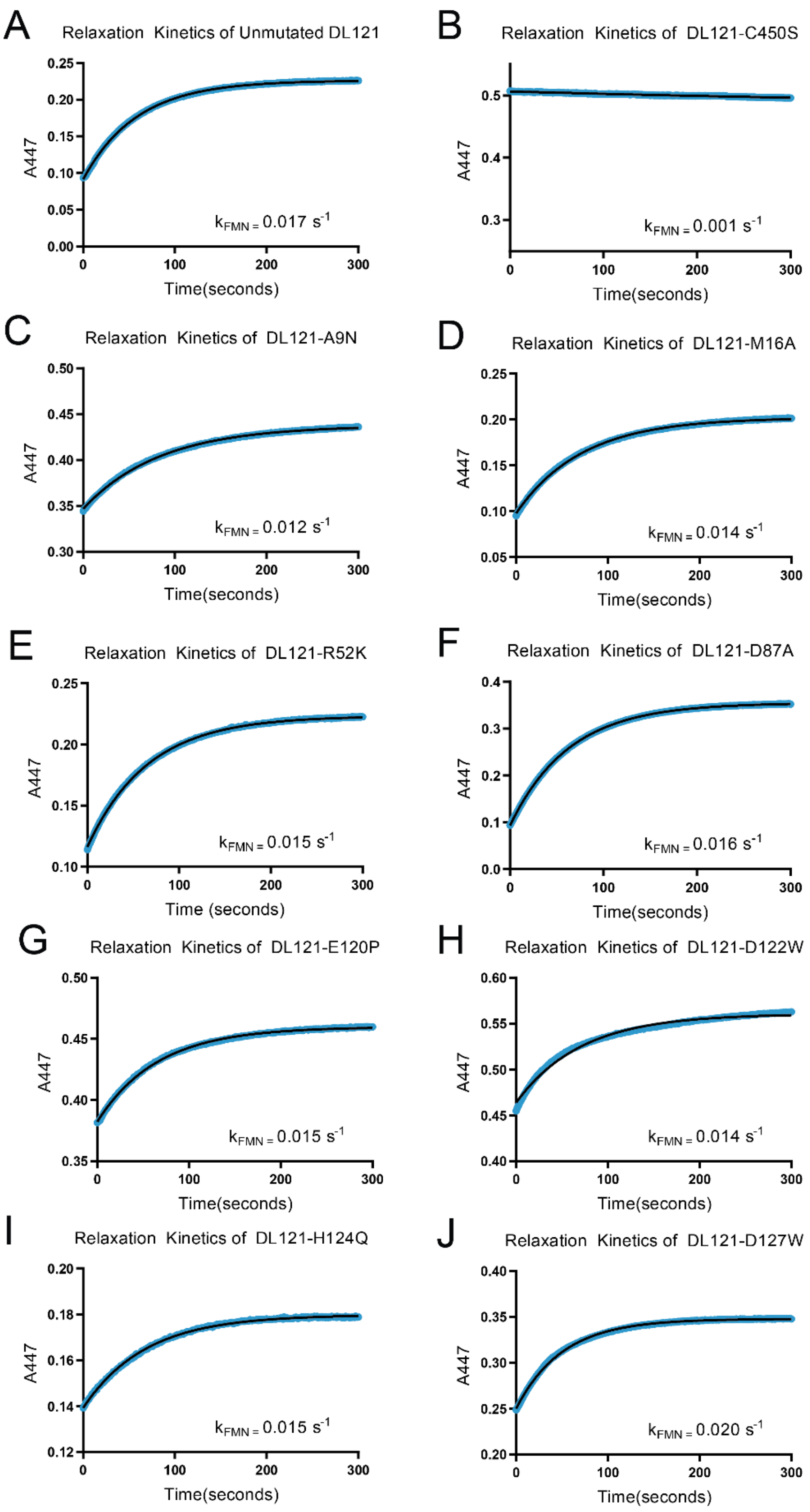
A
Relaxation Kinetics of Unmutated DL121
A447
0.25
0.20
0.15
0.10
0.05
0.00
0
100
200
300
Time(seconds)
$k_{FMN}$ = 0.017 $s^{-1}$
B
Relaxation Kinetics of DL121-C450S
A447
0.5
0.4
0.3
0
100
200
300
Time(seconds)
$k_{FMN}$ = 0.001 $s^{-1}$
C
Relaxation Kinetics of DL121-A9N
A447
0.50
0.45
0.40
0.35
0.30
0
100
200
300
Time(seconds)
$k_{FMN}$ = 0.012 $s^{-1}$
D
Relaxation Kinetics of DL121-M16A
A447
0.25
0.20
0.15
0.10
0.05
0
100
200
300
Time(seconds)
$k_{FMN}$ = 0.014 $s^{-1}$
E
Relaxation Kinetics of DL121-R52K
A447
0.25
0.20
0.15
0.10
0
100
200
300
Time (seconds)
$k_{FMN}$ = 0.015 $s^{-1}$
F
Relaxation Kinetics of DL121-D87A
A447
0.4
0.3
0.2
0.1
0.0
0
100
200
300
Time(seconds)
$k_{FMN}$ = 0.016 $s^{-1}$
G
Relaxation Kinetics of DL121-E120P
A447
0.50
0.45
0.40
0.35
0
100
200
300
Time(seconds)
$k_{FMN}$ = 0.015 $s^{-1}$
H
Relaxation Kinetics of DL121-D122W
A447
0.60
0.55
0.50
0.45
0.40
0
100
200
300
Time(seconds)
$k_{FMN}$ = 0.014 $s^{-1}$
I
Relaxation Kinetics of DL121-H124Q
A447
0.20
0.18
0.16
0.14
0.12
0
100
200
300
Time(seconds)
$k_{FMN}$ = 0.015 $s^{-1}$
J
Relaxation Kinetics of DL121-D127W
A447
0.40
0.35
0.30
0.25
0.20
0
100
200
300
Time(seconds)
$k_{FMN}$ = 0.020 $s^{-1}$

**Figure S10 (previous page):** Relaxation rate of the LOV2 chromophore for select DL121 mutants. The relaxation of the chromophore at 447 nm was observed for 5 minutes following illumination for at least 2 minutes by full spectrum 125 watt 6400K fluorescent lamp (Hydrofarm Inc). With the exception of the allosterically inactivated DL121-C450S all of the assayed LOV2 domains had exponential and reversible relaxation to the dark state near that of the unmutated DL121 ($k_{FMN}$ = 0.017 $s^{-1}$), indicating an active light response in the protein.

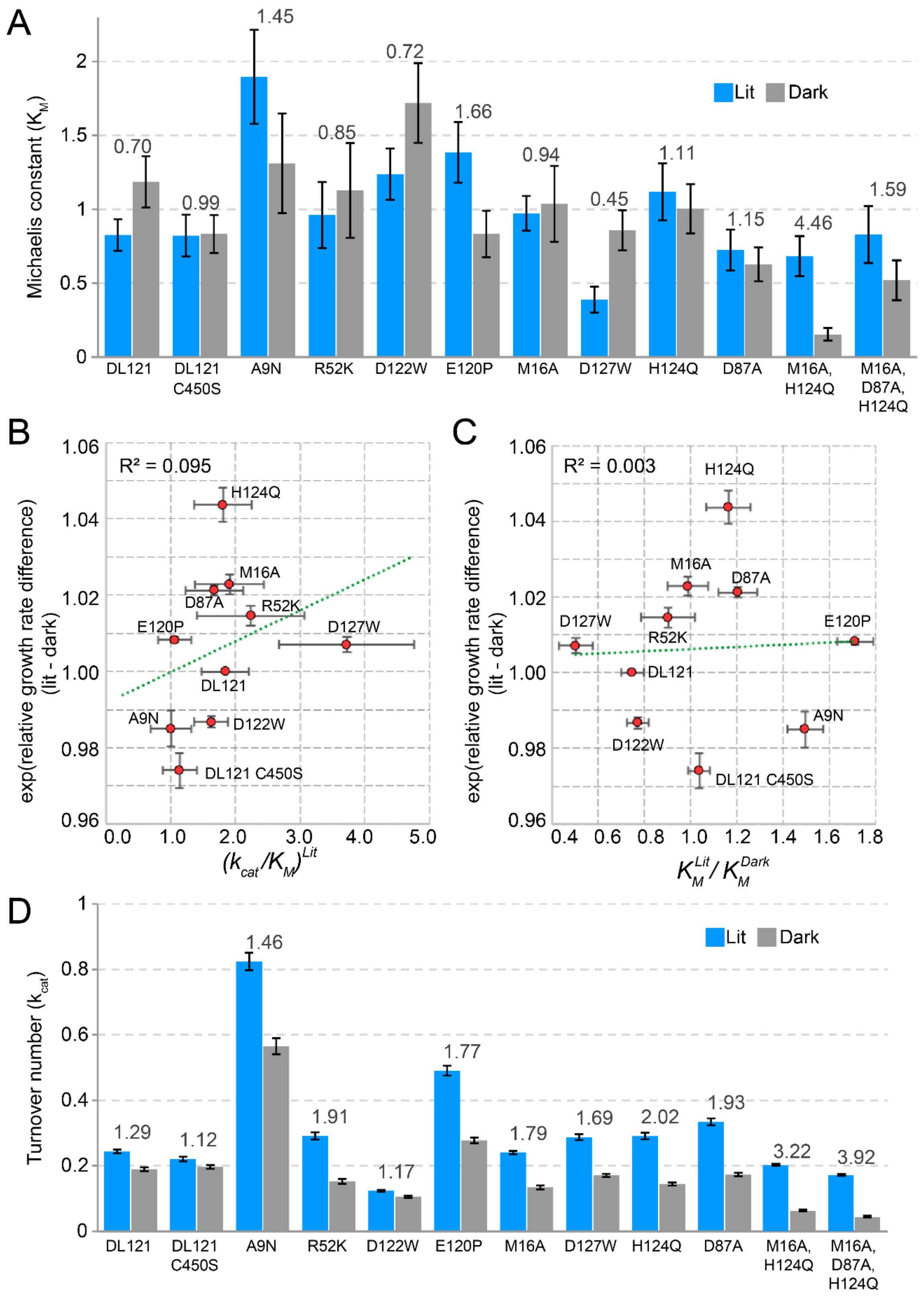
A
Michaelis constant ($K_M$)
Lit
Dark
0.70
0.99
1.45
0.85
0.72
1.66
0.94
0.45
1.11
1.15
4.46
1.59
DL121
DL121 C450S
A9N
R52K
D122W
E120P
M16A
D127W
H124Q
D87A
M16A, H124Q
M16A, D87A, H124Q
B
$R^2 = 0.095$
exp(relative growth rate difference) (lit - dark)
$(k_{cat}/K_M)^{Lit}$
H124Q
M16A
D87A
R52K
E120P
D127W
DL121
A9N
D122W
DL121 C450S
C
$R^2 = 0.003$
exp(relative growth rate difference) (lit - dark)
$K_M^{Lit}/K_M^{Dark}$
H124Q
M16A
D87A
D127W
R52K
E120P
DL121
D122W
A9N
DL121 C450S
D
Turnover number ($k_{cat}$)
Lit
Dark
1.29
1.12
1.46
1.91
1.17
1.77
1.79
1.69
2.02
1.93
3.22
3.92
DL121
DL121 C450S
A9N
R52K
D122W
E120P
M16A
D127W
H124Q
D87A
M16A, H124Q
M16A, D87A, H124Q

**Figure S11 (previous page):** Correlation between *in vivo* allostery and *in vitro* steady state kinetics parameters for mutants of the DL121 fusion. A) The $K_M$ values in the light (blue) and dark (grey) are shown with error bars representing standard error across three replicates. Above each pair of bars the ratio of the lit/dark $K_M$ is shown. The relationship between the relative growth rate difference (*in vivo)* and the ratio of B) catalytic efficiency ($k_{cat}/K_M$) or C) the Michaelis constant ($K_M$). As we expect a log-linear relationship, we compare the ratio of catalytic constants to the exponent of the relative growth rate difference. The green dashed line is the linear regression with the coefficient of correlation ($R^2$) shown. The low coefficient of correlation in both comparisons indicates that there is little relationship between the allosteric growth rate difference and both catalytic efficiency and the Michaelis constant ratios. The error bars represent standard error. D) The turnover number ($k_{cat}$) values in the light (blue) and dark (grey) are shown with error bars representing standard error. Above each pair of bars the ratio of the lit/dark $k_{cat}$ is shown. The Michaelis-Menten kinetics values are reported in Table S1.

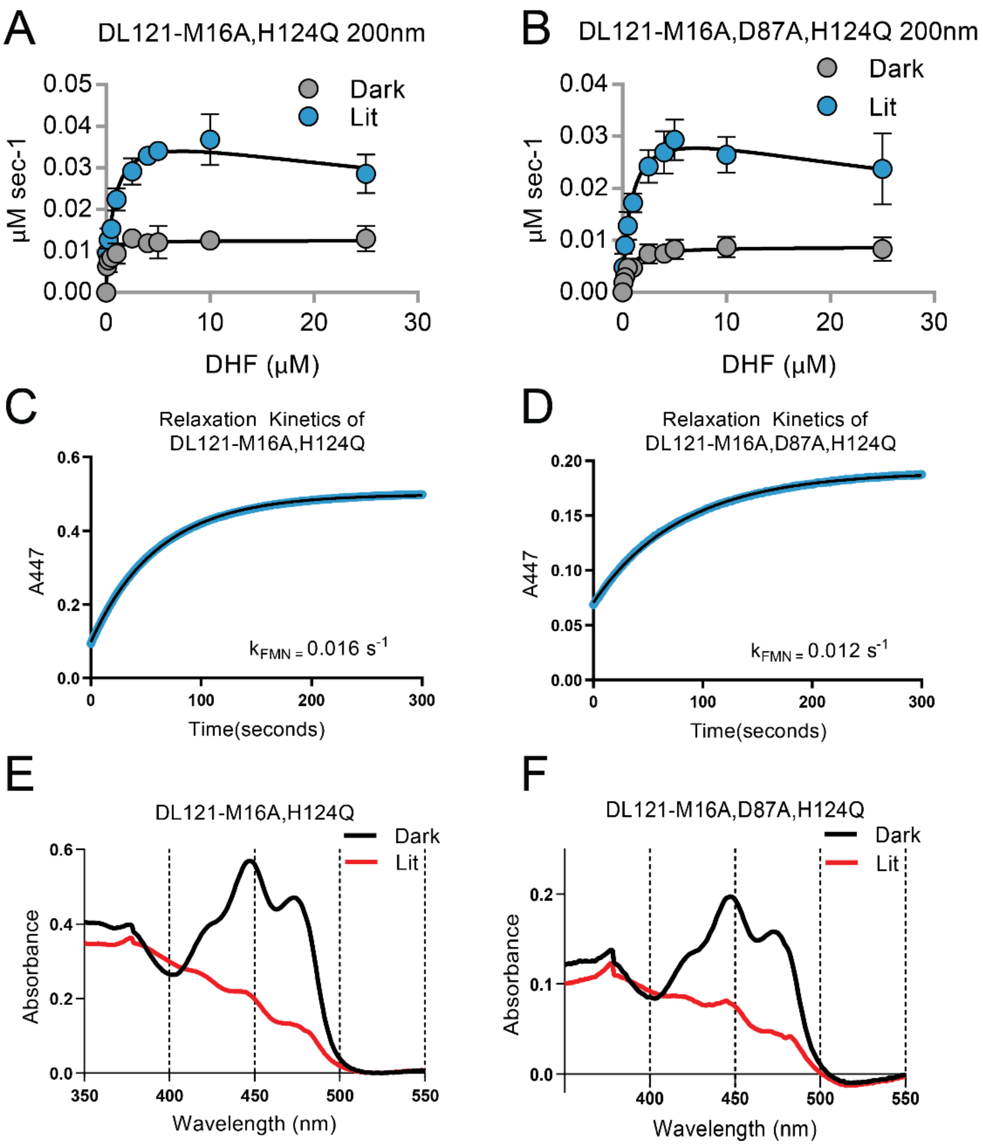


**Figure S12:** Characterization of the DL121- M16A,H124Q and DL121- M16A, D87A, H124Q mutants. A-B) Steady state kinetics measurements in the light and dark. Initial velocity vs. substrate (dihydrofolate) concentration is plotted, Lit (blue) and dark (grey) conditions are shown with error bars representing standard deviation across three replicates. For both the double and triple mutant, the lit states were better fit by a substrate inhibition model than a standard Michaelis-Menten model ($P < 0.05$). The $k_{cat}$, $K_M$, catalytic efficiency and associated error are reported in Table S1. C-D) Relaxation rate of the LOV2 chromophore. The relaxation of the chromophore at 447 nm was observed for 5 minutes following illumination for at least 2 minutes by full spectrum 125 watt 6400K fluorescent lamp (Hydrofarm Inc). E-F) Spectroscopic characterization of LOV2 activation. Lit state absorbance spectra (red line) were measured after illumination for at least 2 minutes by full spectrum 125 watt 6400K fluorescent lamp (Hydrofarm Inc). Dark conditions are taken under the same conditions but using opaque tubes when the sample was placed under the lamp. Both mutants show a characteristic spectral shift upon light stimulation consistent with an active LOV2 domain.

| Mutant | $K_{cat}$ lit | (+/-) | $k_{cat}$ dark | (+/-) | $K_m$ lit | (+/-) | $K_m$ dark | (+/-) | $k_{cat}/K_m$ lit | (+/-) | $k_{cat}/K_m$ dark | (+/-) |
|---|---|---|---|---|---|---|---|---|---|---|---|---|
| Unmutated DL121 | 0.244 | 0.006 | 0.190 | 0.006 | 0.826 | 0.107 | 1.186 | 0.174 | 0.295 | 0.039 | 0.160 | 0.024 |
| DL121 C450S | 0.221 | 0.007 | 0.197 | 0.006 | 0.822 | 0.141 | 0.833 | 0.128 | 0.269 | 0.047 | 0.236 | 0.037 |
| DL121 A9N | 0.824 | 0.027 | 0.566 | 0.024 | 1.896 | 0.318 | 1.311 | 0.337 | 0.435 | 0.074 | 0.432 | 0.113 |
| DL121 M16A | 0.241 | 0.005 | 0.134 | 0.006 | 0.972 | 0.118 | 1.036 | 0.258 | 0.247 | 0.030 | 0.129 | 0.033 |
| DL121 R52K | 0.291 | 0.011 | 0.153 | 0.007 | 0.961 | 0.224 | 1.127 | 0.321 | 0.303 | 0.071 | 0.135 | 0.039 |
| DL121 D87A | 0.334 | 0.011 | 0.173 | 0.005 | 0.724 | 0.138 | 0.627 | 0.115 | 0.462 | 0.089 | 0.276 | 0.051 |
| DL121 E120P | 0.491 | 0.015 | 0.278 | 0.009 | 1.385 | 0.205 | 0.833 | 0.157 | 0.354 | 0.054 | 0.333 | 0.064 |
| DL121 D122W | 0.124 | 0.003 | 0.106 | 0.003 | 1.238 | 0.174 | 1.719 | 0.269 | 0.100 | 0.014 | 0.061 | 0.010 |
| DL121 H124Q | 0.291 | 0.010 | 0.144 | 0.005 | 1.118 | 0.193 | 1.003 | 0.167 | 0.260 | 0.046 | 0.144 | 0.024 |
| DL121 D127W | 0.288 | 0.009 | 0.171 | 0.005 | 0.388 | 0.088 | 0.857 | 0.136 | 0.741 | 0.170 | 0.199 | 0.032 |
| DL121 M16A,H124Q | 0.202 | 0.003 | 0.063 | 0.003 | 0.683 | 0.136 | 0.153 | 0.043 | 0.295 | 0.059 | 0.409 | 0.116 |
| DL121-M16A,D87A,H124Q | 0.171 | 0.003 | 0.044 | 0.002 | 0.828 | 0.193 | 0.519 | 0.135 | 0.207 | 0.048 | 0.084 | 0.022 |

**Table S1**: Steady state kinetic parameters for select point mutants of the DL121 fusion. The parameter $k_{cat}$ is reported in units of $s^{-1}$, $K_m$ is in units of μM. Error is calculated as standard error of the mean over three replicates. Related to Figure 4 of the main text.

| Cutoff for sector definition: | 0.005 | 0.008 | 0.01 | 0.015 |
|---|---|---|---|---|
| Inactivating mutations in sector | 223 | 332 | 373 | 448 |
| Expected by chance | 177 | 280 | 311 | 381 |
| p-value: | $5.73 \times 10^{-7}$ | $2.34 \times 10^{-6}$ | $7.88 \times 10^{-8}$ | $3.19 \times 10^{-8}$ |

**Table S2:** Fisher Exact Test p-values for the null hypothesis that the sector and inactivating mutants are independent properties. Inactivating mutations are defined as those that yield relative growth rates at or below the growth rate for DL121-D27N. Over a range of sector definitions, the null hypothesis is rejected at a confidence level of 0.05 or better, shown in red. Sector definitions were taken from Reynolds et al 2011 [21].

| Cutoff for Conserved Positions: | 1.89 | 1.54 | 1.49 | 1.38 |
|---|---|---|---|---|
| Conserved inactivating mutations | 249 | 377 | 413 | 494 |
| Expected by chance | 175 | 276 | 309 | 374 |
| p-value: | $7.26 \times 10^{-16}$ | $3.78 \times 10^{-20}$ | $8.70 \times 10^{-20}$ | $6.60 \times 10^{-23}$ |

**Table S3**: Fisher Exact Test p-values for the null hypothesis that conserved positions and inactivating mutants are independent. Calculations were made over a range of conservation definitions chosen to result in an equal number positions as the sector positions in Table S2 (23, 36, 40,and 49 positions respectively. In all cases, the null hypothesis is rejected at a confidence level of 0.05 or better (red), and inactivating mutations are enriched at conserved positions beyond expectation due to random chance. Conservation values are calculated as in Reynolds 2011 [21], and reflect the Kullback-Leibler relative entropy of amino acid frequencies at each DHFR position.

| | Sector Cutoff: | 0.005 | 0.008 | 0.01 | 0.015 |
|---|---|---|---|---|---|
| **Cutoff for allostery enhancing:** | Allosteric positions in sector: | 4 | 5 | 5 | 7 |
| | Expected | 13 | 22 | 25 | 31 |
| $p < 0.05$ | p-value: | 0.008 | $3.43 \times 10^{-5}$ | $5.82 \times 10^{-6}$ | $3.52 \times 10^{-7}$ |
| $p < 0.016$ | # in sector: | 0 | 0 | 0 | 1 |
| | Expected | 6 | 11 | 12 | 15 |
| | p-value: | 0.013 | $4.23 \times 10^{-4}$ | $1.68 \times 10^{-4}$ | $4.64 \times 10^{-5}$ |

| | Sector Cutoff: | 0.005 | 0.008 | 0.01 | 0.015 |
|---|---|---|---|---|---|
| **Cutoff for allostery disrupting:** | Allosteric positions in sector: | 2 | 15 | 15 | 15 |
| | Expected | 5 | 8 | 9 | 11 |
| $p < 0.05$ | p-value: | 0.257 | 0.013 | 0.038 | 0.25 |
| $p < 0.016$ | # in sector: | 1 | 7 | 7 | 7 |
| | Expected | 1 | 2 | 3 | 3 |
| | p-value: | 0.967 | 0.004 | 0.010 | 0.049 |

**Table S4:** Fisher Exact Test p-values for the null hypothesis that the sector and allosteric mutations are independent. We compared over four sector cutoffs (as defined in [21]) and at two cutoffs for allostery significance (a standard p-value of 0.05, and an adjusted p-value of 0.016). The multiple hypothesis testing adjusted p-value was obtained by Sequential Goodness of Fit (SGoF, [49]). The top table shows the association between sector positions and allostery enhancing mutations; the bottom table computes the associate between sector positions and allostery disrupting mutations. In nearly all cases, the null hypothesis is rejected at a confidence level of 0.05 or better, shown in red.

| | Allosteric Surface Positions | |
|---|---|---|
| **Cutoff for allostery enhancing:** p < 0.05 | Observed | 88 |
| | Expected | 73 |
| | p-value: | 0.004 |
| p < 0.016 | Observed | 44 |
| | Expected | 35 |
| | p-value: | 0.013 |

| | Allosteric Surface Positions | |
|---|---|---|
| **Cutoff for allostery disrupting:** p < 0.05 | Observed | 33 |
| | Expected | 27 |
| | p-value: | 0.061 |
| p < 0.016 | Observed | 12 |
| | Expected | 8 |
| | p-value: | 0.048 |

**Table S5:** Fisher Exact Test p-values for the null hypothesis that the solvent accessible DHFR surface and allosteric mutations are independent. At two cutoffs for allostery (a standard p-value of 0.05, and an adjusted p-value of 0.016), the null hypothesis is rejected at a confidence level of 0.05 or better, shown in red.

| | Sector Cutoff: | 0.005 | 0.008 | 0.01 | 0.015 |
|---|---|---|---|---|---|
| **Cutoff for allostery enhancing:** | Allosteric surface positions within or connected to sector: | 2 | 13 | 15 | 28 |
| | Expected | 18 | 27 | 28 | 33 |
| $p < 0.05$ | p-value: | $4.88 \times 10^{-5}$ | 0.002 | 0.006 | 0.360 |
| | # connected: | 10 | 5 | 7 | 15 |
| $p < 0.016$ | Expected | 1 | 13 | 13 | 16 |
| | p-value: | 0.009 | 0.019 | 0.074 | 0.987 |

| | Sector Cutoff: | 0.005 | 0.008 | 0.01 | 0.015 |
|---|---|---|---|---|---|
| **Cutoff for allostery disrupting:** | Allosteric surface positions within or connected to sector: | 19 | 22 | 22 | 24 |
| | Expected | 6 | 10 | 10 | 12 |
| $p < 0.05$ | p-value: | $1.56 \times 10^{-7}$ | $1.52 \times 10^{-5}$. | $2.72 \times 10^{-5}$. | $6.60 \times 10^{-5}$ |
| | # connected: | 8 | 9 | 9 | 10 |
| $p < 0.016$ | Expected | 2 | 3 | 3 | 4 |
| | p-value: | $1.38 \times 10^{-5}$ | $2.30 \times 10^{-4}$ | $3.16 \times 10^{-4}$ | $2.50 \times 10^{-4}$ |

**Table S6:** Statistical association of allosteric mutations and surface positions that are either within or contacting the sector. Contacting was defined as two atoms within the sum of their Pauling radii plus 20%. A surface site contacts the sector if the peptide bond atoms of the surface site contact any atoms in the sector position. P-values were computed by Fisher exact test with the null hypothesis that the sector and allosteric mutations are independent. Cutoffs for sector definition as defined in [21] are shown as well as mutants determined to effect allostery either at a 95% confidence interval ($p < 0.05$) or at the multiple hypothesis testing adjusted p-value ($p < 0.016$). The null hypothesis that there is no relationship between allosteric mutations and sector or sector contacting positions on the surface of DHFR of the DL121 chimera is rejected at a confidence level of 0.05 or better over a range of cutoffs, shown in red. Allostery enhancing mutations are depleted from sector connected surface sites, while allostery disrupting mutations are enriched (in comparison with random expectation).

| | Sector Cutoff: | 0.005 | 0.008 | 0.01 | 0.015 |
|---|---|---|---|---|---|
| **Cutoff for allostery enhancing:** | Allosteric surface positions connected to sector: | 2 | 13 | 15 | 28 |
| | Expected | 13 | 18 | 17 | 19 |
| p < 0.05 | p-value: | 0.001 | 0.269 | 0.697 | 0.033 |
| | # connected: | 1 | 5 | 7 | 15 |
| p < 0.016 | Expected | 6 | 8 | 8 | 9 |
| | p-value: | 0.041 | 0.275 | 0.837 | 0.049 |

| | Sector Cutoff: | 0.005 | 0.008 | 0.01 | 0.015 |
|---|---|---|---|---|---|
| **Cutoff for allostery disrupting:** | Allosteric surface positions connected to sector: | 18 | 7 | 7 | 9 |
| | Expected | 5 | 6 | 6 | 7 |
| p < 0.05 | p-value: | $3.83 \times 10^{-10}$ | 0.971 | 0.883 | 0.379 |
| | # connected: | 8 | 2 | 2 | 3 |
| p < 0.016 | Expected | 1 | 2 | 2 | 2 |
| | p-value: | $8.02 \times 10^{-8}$ | 0.731 | 0.772 | 0.777 |

**Table S7:** Statistical association of allosteric mutations and surface positions that are contacting the sector. In contrast to Table S6, surface positions within the sector are excluded. Fisher Exact Test p-values were calculated for the null hypothesis that the sector and allosteric mutations are independent. Cutoffs for sector definition as defined in [21] are shown as well as mutants determined to effect allostery either at a 95% confidence interval ($p < 0.05$) or at the multiple hypothesis testing adjusted p-value ($p < 0.016$). At most cutoff combinations, there is not a statistically significant association between sector connected surface sites and either mutations that enhance (top panel) or disrupt allostery (bottom panel).
